# Kintsugi decides, gene by gene, where spatial transcriptomics borrows information

**DOI:** 10.64898/2026.08.30.748061

**Authors:** Chen Yang, Xianyang Zhang, Jun Chen

## Abstract

Subcellular spatial transcriptomics captures where RNA is in tissue, but a single location holds too few molecules of any one gene to estimate composition alone. Every current method fixes in advance where to borrow — a smoothing scale, a cell outline or a factor model — and the fixed choice shapes what is visible. Kintsugi removes the fixed choice and lets held-out molecules decide, gene by gene, how much to borrow from spatial neighbours and from other genes at the same location. On a lung section measured by both Xenium and Visium HD, the data-chosen allocation placed an epithelial programme where the Xenium molecules were, ahead of smoothing, cell segmentation and a factor model; the result replicated across tissues and against protein. Across a 45-core pulmonary fibrosis cohort, separating composition from captured amount shows that a fibroblastic focus is not a place with more RNA but a place with different RNA: 2.8-fold higher in activated-fibroblast composition while segmented nuclear density is at most 1.08-fold higher.

## 1 Introduction

Subcellular spatial transcriptomics measures both the amount of RNA captured from a location and the identities of the captured molecules. Platforms such as Visium HD[1] and Stereo-seq[2] now resolve tissue below or near cellular dimensions, where tissue organisation extends beyond detected nuclei: neuronal processes carry spatially localised transcripts whose compositions differ from the cell body[3, 4]. Interpreting these measurements requires keeping every spatial location, including those without an assigned cell, and distinguishing measured counts from estimated expression.

Existing approaches each fix the source of borrowed information. Spatial methods pool within a chosen scale — fixed grids or smoothing kernels — and changing the scale changes the summary[5–7]. Cell-assignment methods (Bin2cell[8], BIDCell[9], ENACT[10], Cellist[11]) borrow within an inferred cell outline. FICTURE[12] borrows across genes through local factor mixtures at a fixed rank and anchor scale. Reference-based deconvolution (cell2location[13]) and reference-based annotation (TopACT[14]) borrow from an external single-cell atlas. Each fixes the allocation in advance; none lets the data choose, per gene, how much of each source to use. A representation can retain all tissue locations yet pool them at a scale that obscures a particular biological contrast; conversely, useful local predictions need not define regions that correspond to individual cells.

RNA amount and gene composition pose different statistical problems. Total captured counts, summed from hundreds of molecules in a typical bin, can be estimated at the native resolution. Individual gene proportions, resting on far fewer counts per gene, require borrowing from neighbours — and the useful neighbourhood depends on the gene, its depth and whether a compositional boundary lies nearby. A spatial neighbourhood that estimates the total reliably may therefore contain too little evidence for individual genes, while a larger neighbourhood can average across compositional boundaries. Requiring both quantities to share one resolution makes this trade-off implicit; keeping them separate makes each target explicit.

Here we let held-out molecules resolve that trade-off. How much to borrow for one gene at one location depends on the gene, on the scale of the programme it belongs to and on whether a compositional boundary lies nearby. Kintsugi splits captured molecules into disjoint sets and lets held-out molecules decide, for every gene separately, how much to borrow from spatial neighbours — connected regions whose boundaries track compositional changes — and from a factor component fitted on the same regions. Density is estimated separately from the native totals, which need no allocation across genes. We test the fields where they can be falsified: against Xenium molecules on the same section as Visium HD, against protein channels on the same Xenium section, and against pathologist annotations across a pulmonary fibrosis cohort, where separating composition from captured amount changes the reading of the fibroblastic focus, the lesion that defines disease activity in IPF.

## 2 Results

### 2.1 Two fields, and a decision about where each gene borrows information

Kintsugi represents two properties of the observed RNA field: captured density, *ρ_i_*, in molecules per *µ*m^2^, and conditional gene composition, *π_ig_*, the probability of a gene among the molecules captured at bin *i*. Their product is the captured intensity of gene *g* per unit area (Fig. 1a,b). The Poisson count likelihood factors into a total-count term and a gene-allocation term, so the two are estimated separately (Methods). Density is estimated from the totals with one validation-selected pooling, and validation asks for almost none of it: 506 of 510 fits across the six datasets of Fig. 1d chose the native bin or a half-bin kernel, because a bin’s total is already measured by all of its molecules. Composition is where the difficulty lies: a bin holds too few molecules of any one gene, so its composition must be borrowed, and the question is from where. Every fit borrowed for composition and almost none for density, which is the practical reason to estimate the two separately; what a fit borrowed from varied by dataset, from about half the weight on a gene’s own spatial pooling in the lung microarray to less than one per cent in developing cartilage, averaged over fits.

**Fig. 1.**
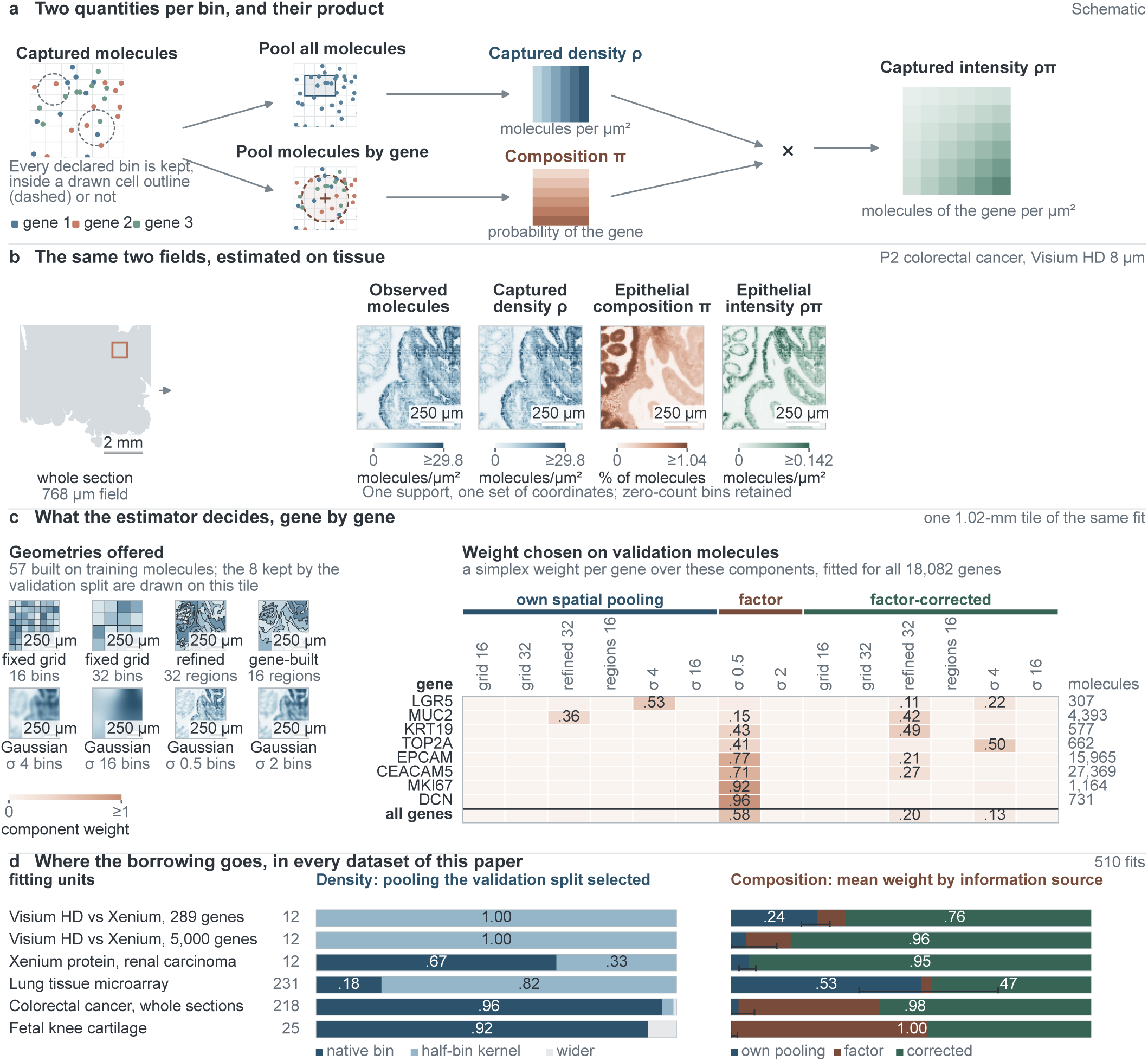
Two estimated fields on a common tissue support, and where each gene borrows its composition. **a**, Schematic. Every native bin carries two estimates: captured density *ρ*, pooled from all molecules, and conditional composition *π*, the probability of a gene among the molecules captured there, pooled gene by gene. Their product is that gene’s captured intensity per unit area. Every declared bin is kept, inside a drawn cell outline or not. **b**, The same quantities on colorectal cancer section P2 (Visium HD, 8-*µ*m bins, 545,913 support bins, 18,082 genes)[1], fitted whole with the tiled estimator, molecules split 50/25/25 into training, validation and test counts (seed 115,000). The 768-*µ*m field is exactly the 9,216 bins owned by the single tile of **c**, so both panels describe one model. Observed molecules and density share a colour maximum; composition and intensity use their own, higher values saturated. Density reproduces the observed totals (Pearson *r* = 0.9995, median ratio 1.00) because the validation counts chose no pooling for density, the general outcome quantified in **d**; composition is where the borrowing happens. The two fields are not redundant: bins in the top decile of epithelial composition hold fewer molecules than the lower half (377 against 654 per bin), and *ρ* and *π* rank bins differently (Spearman −0.11). The epithelial programme is EPCAM, KRT8 and KRT19; KRT18 is absent from the 18,082-symbol universe shared by all three colorectal sections. **c**, What the fit decided inside the 1.02-mm tile holding the field of **b**. Left, the 57 geometries the bank built from training counts alone (6 fixed grids, 45 partitions built from all genes by joint Poisson merge and perimeter-penalised refinement, 6 Gaussian kernels); the 8 drawn are those validation kept. Right, the simplex weight fitted on validation counts for every gene over the 14 selected components, those geometries taken three ways: the gene’s own pooling, a factor component, and the factor prediction corrected by that pooling. Eight named genes span the depth range, with their training molecules. The last row, the mean over all 18,082 genes, places 0.58 on the factor component, 0.20 and 0.13 on two factor-corrected components and 0.078 in total on the genes’ own pooling, so the named genes are the tails of that distribution and not selected extremes. Rows sum to one and weights below 0.10 are unlabelled. Regions are not cells and weights are not cell types. **d**, The same decision across every dataset, over 510 fits: 12 window fits for each of three window studies, 231 microarray tiles, 218 colorectal tiles and 25 cartilage tiles. Left, the share of fits selecting the native bin, a half-bin Gaussian kernel or anything wider for density; 506 of 510 took one of the first two, so density needs almost no pooling anywhere. Right, the mean composition weight staying with the gene’s own pooling against the mean going to factor-informed components; the foot bar is the 10th to 90th percentile of the own share. The factor and factor-corrected sub-bands are drawn but not quantified: they trade against one another, the colorectal median factor weight running from 0.70 in P2 to 0.00 in P5, while their sum stays between 0.97 and 0.99 in all three sections. How much stays with the gene is a property of the dataset, from 0.53 in the microarray to 0.005 in cartilage; no fit anywhere used the section-wide spectrum. These are fit-level mixture weights, not the per-gene matrix of **c**; on this tile the two agree to within 0.07 in every component. Bilous NSCLC is absent because its records carry no final block, not by choice.

Molecules are split by binomial thinning into training, validation and test counts. Training counts supply the two kinds of neighbour. Spatial neighbours are defined by connected regions built from all genes at once by a joint Poisson merge tree whose edges are then moved, under a perimeter penalty, to where composition changes; fixed grids and Gaussian kernels are offered alongside, each at a fine and a coarse scale. Molecular neighbours are defined by a factor model fitted on training counts, evaluated on the same regions, which lets a sparse gene take its local level from the genes it co-varies with, and by corrected components in which the factor prediction is the shrinkage centre that a gene’s own spatial pooling overrides in proportion to the molecules the chosen neighbourhood holds at that bin. Validation counts choose the region count, the edge penalty, the scales and the factor rank, and fit for every gene a simplex weight over these sources, renormalised within each bin. On real panels almost every gene put most of its weight on the factor-informed components, and the global spectrum never entered the mixture, having lost to a pooled geometry in every family. Within a corrected component the share of a gene’s own pooling that survives is set by the bin’s molecules and the selected shrinkage rather than by the gene. Factor-informed components dominate on both panels, and with 5,000 genes they alone reproduce the complete field on every endpoint. The weights themselves are a fitting device rather than a measurement of where a gene borrows: the validation objective is nearly flat between the factor component and its corrected siblings, so the split between them moves with the optimiser’s starting point and with the shrinkage menu while the estimated composition does not (Supplementary Note 9). We therefore report the field and its held-out scores rather than per-gene source attributions. Test counts are scored only. The resulting probabilities are defined on the complete declared tissue support, including bins with no observed molecules and molecules assigned to no cell; neither regions nor weights are interpreted as cells or cell-type proportions (Fig. 1c).

Published CRC singlet annotations illustrate why complete support is retained: restriction to singlets changed the represented tumour, stromal and immune fractions across three patients[1] (Extended Data Fig. 1). Those calls were derived from expression deconvolution; the field estimators retain declared tissue bins without requiring a confident cell-type assignment.

Held-out molecules of the same modality test whether an estimator predicts, not whether it places a programme where it belongs. We therefore evaluate placement against references that never enter the fit: known truth, Xenium molecules on the same section as Visium HD, protein channels on the same Xenium section, and pathologist annotations across a cohort, where separating composition from captured amount changes the reading of the fibroblastic focus. We ablate the estimator by information source rather than by implementation.

### 2.2 Spatial neighbours must be defined by the genes, and the decision must be per gene

We first asked, with known truth, what spatial borrowing requires. On a 32 × 32 lattice with 16 genes, a three-gene programme and Poisson counts at 10 or 80 molecules per bin (20 draws each), four scenarios placed the programme in a smooth field (negative control), in a step of composition at constant density, in a step aligned with a density step, or in a narrow band five bins wide (Methods). Every estimator used the same thinning, candidate menus and selection protocol; a separately tuned Gaussian, a Gaussian tuned for the programme alone (target-tuned) and an oracle that pools within the true regions bounded the comparison.

Geometry built from total counts cannot see a composition-only step: the previous total-count guide and a greedy merge of total counts gave the same programme edge error as each other (mean absolute error at the boundary, 0.0273). The same merge algorithm applied to the full gene cube reduced the error to 0.0234 in the mixed field and to 0.0115 for the adaptive component alone, better than the target-tuned Gaussian (0.0227) in 20 of 20 draws (Fig. 2a). The gain is attributable to composition information in the partition, because the algorithm is otherwise identical.

**Fig. 2.**
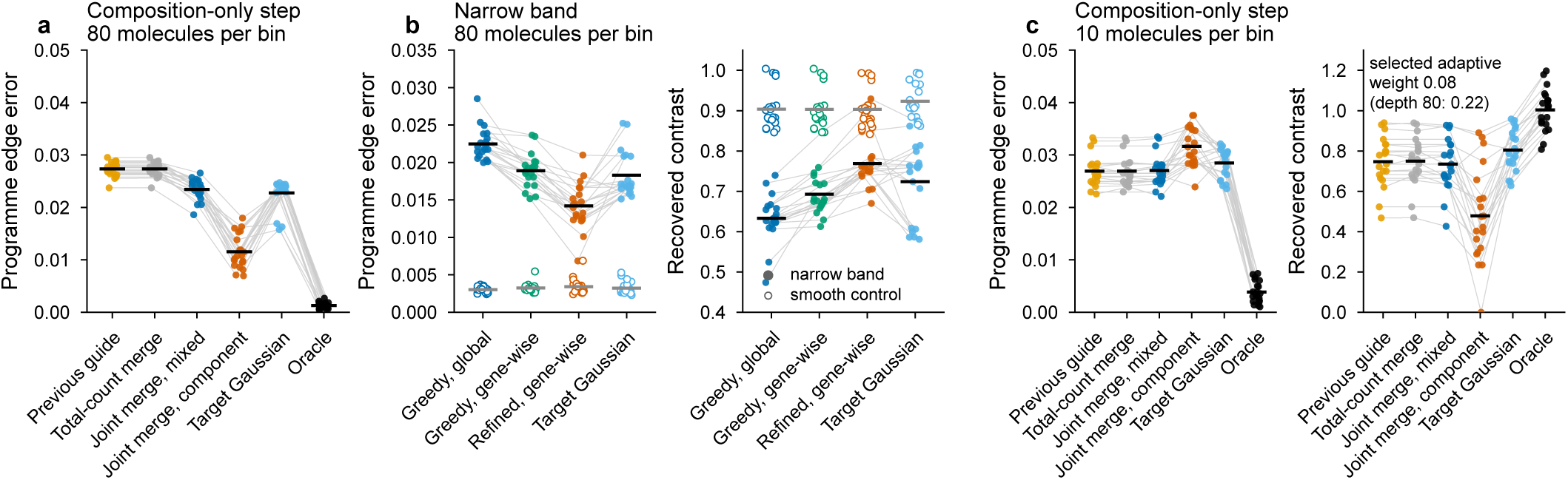
Spatial neighbours must be defined by the genes, and the decision must be per gene. **a**, Known-truth composition-only step at 80 molecules per bin: programme edge error for the previous total-count guide, a greedy merge of total counts, the joint multi-gene merge (mixed field and adaptive component alone), the target-tuned Gaussian and the oracle; 20 draws, paired by draw (grey lines), bars are means. **b**, Narrow band: edge error and recovered contrast for greedy merge with global weights, gene-wise weights, and refined partitions with gene-wise weights, against the target-tuned Gaussian; the smooth control is shown for the same estimators (open symbols). **c**, Depth limit: edge error and recovered contrast for the estimators in **a** on the composition-only step at 10 molecules per bin; the validation-selected adaptive weight is annotated.

Two further changes were needed before the all-gene model matched a Gaussian tuned for the programme on the narrow band. With one global weight vector, validation counts held the adaptive component at 0.2–0.3 even where it was the best component for the programme; gene-wise weights raised the programme genes’ adaptive weight to 0.5–0.7 while background genes stayed near 0.25, and improved every endpoint in 20 of 20 draws in every structured scenario, with no change in the smooth control. Boundary refinement then removed the cost of ragged edges (validation chose *λ* = 3 or 10, never 0): the all-gene field reached an edge error of 0.0142 against 0.0183 for the target-tuned Gaussian and a recovered contrast of 0.77 against 0.72 (17 and 19 of 20 draws; Fig. 2b). Each of the three choices was necessary and together they were sufficient here. Molecular borrowing has nothing to offer with sixteen genes, and validation molecules left the factor component 0.04–0.13 of the programme weight and no endpoint moved. What the design requires of an uninformative source is that it change no endpoint; it does not always receive near-zero weight, and a case where it takes weight without changing an endpoint is reported in Supplementary Note 7. The complete estimator improved the all-gene held-out score in 19–20 of 20 draws at 80 molecules per bin but placed the edge slightly worse than the spatial-only fields (0.0135 to 0.0174 on the aligned step; smooth control and 10-molecule depth unchanged): validation rewards the corrected components because they predict better across all sixteen genes, while the sharper spatial components place this one edge better.

With 300 genes and a truth of eight co-expressed programmes, the same estimator selected the molecular source and used it: against a flat background it placed the target edge better than spatial borrowing alone at both depths (edge error 0.025 against 0.030 at 10 molecules per bin, 0.005 against 0.016 at 80; 19–20 of 20 draws), with 0.9 of the target genes’ weight on factor-informed components. Because the factor field’s geometry is chosen once on the evidence of all genes, a programme whose edge is sharper than the panel average is placed less well even as the held-out score improves; the effect is bounded and its dependence on depth and panel size is reported in Extended Data Fig. 2 and Supplementary Note 7.

The simulations also mark where spatial borrowing stops. In this sixteen-gene truth at 10 molecules per bin, borrowing from neighbours recovered less than half of the composition-only step (contrast 0.48) while a Gaussian tuned on the three target genes alone recovered 0.80; sixteen genes leave molecular borrowing nothing to offer, so the validation-selected weight on the factor component stayed at 0.08 and the estimator returned the spatial answer, doing no harm and no good (Fig. 2c). What is exhausted there is the evidence, not the depth. In the operating-range grid (300 genes, no co-expression between the target and other programmes), a three-gene programme stepping over the same range at the same depth was recovered at 0.73 of its true contrast against 0.40 for spatial borrowing alone, and its genes placed 0.97 of their weight on factor-informed components rather than the 0.08 here. Spatial borrowing alone needed between 40 and 80 molecules per bin to reach 0.73, so at that point a molecular source was worth about five-fold depth. Across a grid of depth, boundary contrast and programme size, what decides whether a boundary is recoverable is not the depth of a bin but how many molecules the programme has in the analysis window and how much its share changes across the boundary. Writing *f* for that relative change and *M* for the programme’s molecules in the window, boundaries with *f √M* above about 60 were seen and their amplitude returned, which is *M* of at least about 3,600*/f* ^2^; over 162 grid settings the rule had four exceptions, all of them boundaries just below the threshold that were recovered anyway, and it held unchanged when the bin edge was varied four-fold, over which the per-bin depth changes by sixteen. Inside that recoverable region, and against a flat background, the complete estimator never fell clearly behind a Gaussian tuned on the programme itself; the settings where it did fall behind are those where neither recovers the boundary at all (Extended Data Fig. 3). Where the background itself varies in space, the loss in edge placement described above returns, and the recoverable region marks where a boundary can be seen and its amplitude returned rather than where it is placed best.

### 2.3 Where a programme is placed, by information source, on one section measured twice

The simulation says what the design must contain; whether the data-chosen allocation places a programme where the molecules are is a question only an independent measurement of the same tissue can answer. In each of two experiments, a 5-*µ*m FFPE lung adenocarcinoma section was measured first on Xenium and then on Visium HD[15], using a 289-gene panel in Experiment 1 and a 5,000-gene panel in Experiment 2, of which 282 and 4,043 genes are shared; the 3,000 deepest per window were carried through every estimator. The Xenium molecules, binned on the Visium HD 8-*µ*m grid through the vendor’s alignment of the shared H&E scan and refined by a one-bin shift chosen on total counts only, serve as a positional reference that never enters fitting or selection (Methods). Four 1-mm windows were chosen by a rule on the Xenium epithelial fraction alone, and the epithelial programme (EPCAM, CDH1, MUC1, plus NKX2-1 where the panel carries it) was fixed from panel membership before any fit.

Twelve window seed units score each estimator on the rank agreement of its programme probability with the Xenium fraction in the edge zone, the area under the curve against the Xenium compartment, the area-matched overlap and boundary distance, the held-out Xenium likelihood and the programme contrast relative to raw pooled counts.

We ablate the estimator by what it is allowed to borrow. Spatial neighbours alone already placed the programme ahead of a separately tuned Gaussian in 12 of 12 units (edge-zone Spearman correlation 0.427 against 0.252, AUC 0.792 against 0.666; Fig. 3a,b); offering a factor component raised every endpoint in 12 of 12 (Extended Data Table 1), with almost all of the positional gain attributable to the molecular source: letting each gene correct the factor prediction rather than the global spectrum bought amplitude and held-out likelihood without changing placement. The complete estimator reached 0.510, 0.846, overlap 0.809 and 0.94 of the raw pooled contrast, above both comparators that were told the target in 12 of 12 units (programme versus rest on gene-built regions 0.428, 0.800; the same fit restricted to Gaussian kernels 0.374, 0.768). Validation molecules gave the three epithelial genes 0.80–0.99 of their weight to the factor-informed components; the section-wide spectrum is absent from the mixture because no family’s winner was the unpooled estimate, so it is not a source these genes declined but one they were never offered (Fig. 3). The factor component alone was worse than the combination on the all-gene held-out score, so neither source suffices by itself. Replacing the per-gene weights with a single global weight vector and changing nothing else isolated the per-gene decision. On the 289-gene panel it cost a tenth of the complete estimator’s advantage over raw pooled counts (edge AUC 0.846 to 0.823, overlap 0.809 to 0.788, 12 of 12 units), and the programme genes’ weights lay far from the global vector (total variation 0.52–0.61). On the 5,000-gene panel the per-gene weights sat close to the global vector and left positional endpoints unchanged while still improving held-out likelihood. Deciding per gene is what makes a marker panel’s geometry recoverable; it is not a uniform improvement.

**Fig. 3.**
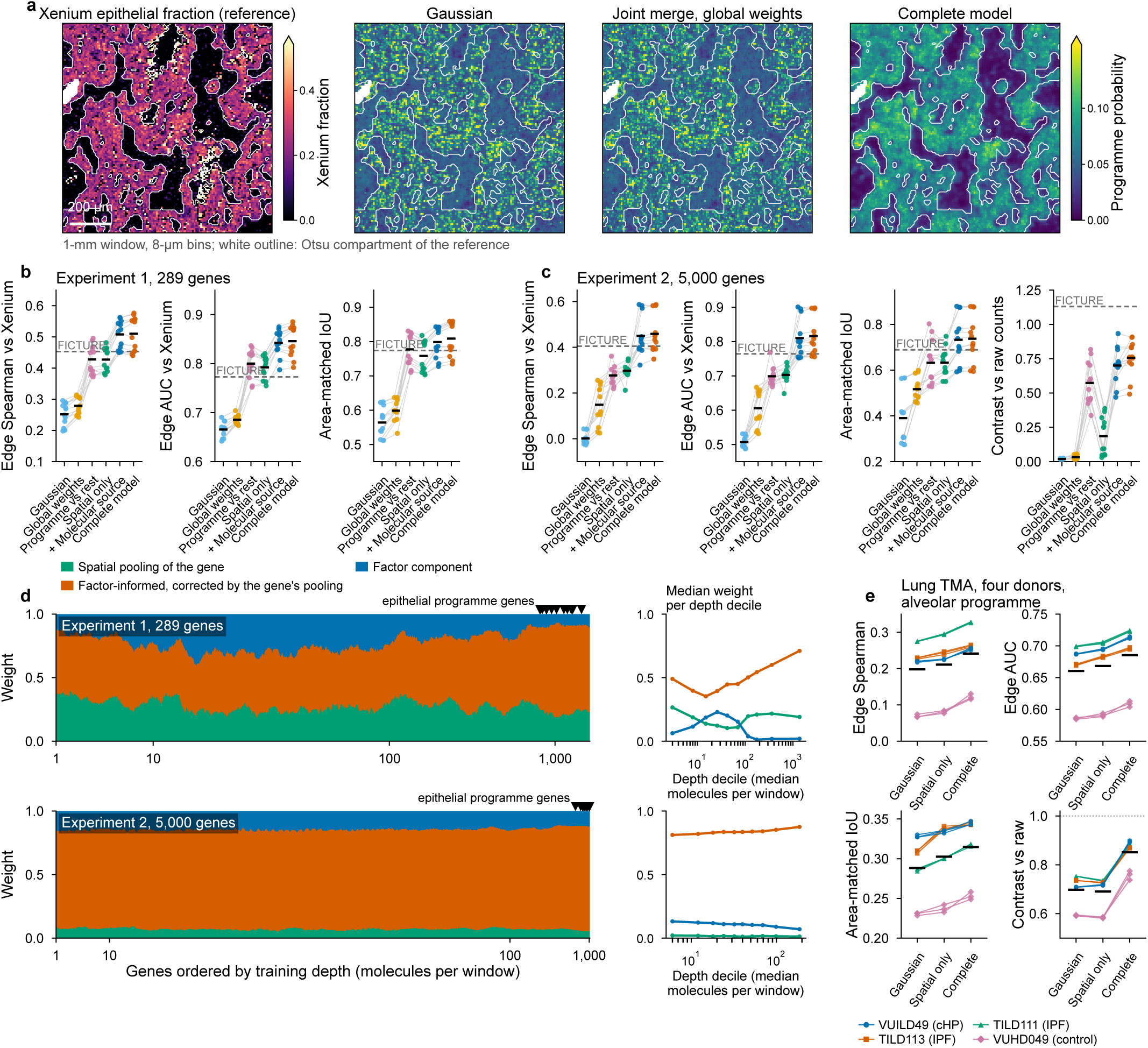
Each gene decides where to borrow, and the programme lands where Xenium finds the molecules. **a**, One 1-mm window of the post-Xenium lung section[15]: Xenium epithelial fraction on the Visium HD 8-*µ*m grid (reference), and the epithelial programme probability from the Gaussian, the joint merge with one global weight vector, and the complete model; the Otsu compartment of the reference is outlined in white. **b**, Experiment 1 (289 genes), 12 window × seed units: edge-zone Spearman correlation with the Xenium fraction, edge AUC and area-matched intersection over union per estimator. “Spatial only” is the estimator restricted to spatial neighbours (regions built from the genes at two scales, per-gene weights); “+ molecular source” adds factor components on every geometry without correction or refit; the programme-versus-rest model is the reference and the dashed line is FICTURE’s best setting given all Visium HD molecules. **c**, Experiment 2 (5,000 genes, 0.23 programme molecules per bin): the same endpoints and the contrast relative to raw pooled counts, showing the amplitude limit of spatial borrowing alone and its recovery by molecular borrowing. **d**, Validation-selected weight of every gene on three source classes (spatial pooling of the gene’s own counts; factor-informed components in which the gene’s spatial pooling corrects the factor prediction; the factor component itself). The section-wide spectrum is not shown because it never enters the mixture: it is each family’s starting value and is displaced by that family’s winner in every window. Genes are ordered by training depth (molecules per window), four windows pooled, one seed; stacked areas are running means over about one thirtieth of the genes, and triangles mark the epithelial programme genes. Right, the median weight of each class per depth decile. Genes at every depth put most of their weight on factor-informed components. On the 5,000-gene panel the corrected components hold 0.78 to 0.80 of the weight in every decile, with no depth structure and none needed. On the 289-gene panel the factor component is chosen mainly at intermediate depth; that trend is about three times the spread across optimiser restarts, but the deepest decile’s factor weight is the least stable point of the curve and a restart can move it threefold, so the decline at the deep end should not be read as a preference (Supplementary Note 9). Class weight records which component a gene chooses, not how much of its own pooling that component keeps: inside a corrected component that share is *S_i_/*(*S_i_* + *τ* ), a property of the bin and of the selected *τ* , so the corrected class returns about 40% of a gene’s own pooling on the 289-gene panel but about 7% on the 5,000-gene panel, where the selected shrinkage leaves the corrected components within 5% of the factor field itself. That is a property of the parametrisation rather than of the tissue: a corrected component converges on the factor field as its shrinkage grows, so the two classes are near-duplicates and only their sum is determined. **e**, Replication on a lung tissue microarray measured by Xenium and then by Visium HD on the same sections (four donors, one core each, three seeds; 12 core × seed units, alveolar programme): the same endpoints for the Gaussian, the spatial-only fields and the complete model; lines join the same unit, colour and symbol mark the donor, black bars are means (Extended Data Fig. 4 shows the maps). Units are technical replicates of one section per experiment in **b**–**d** and of four donor sections in **e**; the Xenium reference never enters fitting or selection.

The two fixed allocations in use sit on either side of this space, and both were run on the same section and scored on the same windows (Extended Data Table 1). FICTURE[12] borrows across genes through local factor mixtures. We evaluated fixed rank and spatial settings without held-out selection; run with its authors’ implementation over six settings, it was below the spatial-only fields on AUC, overlap and Xenium likelihood in 4 of 4 windows when trained on the same molecules, and with all Visium HD molecules and the setting chosen on the endpoint itself it reached the level of the spatial-only fields on rank agreement (0.453 against 0.427) but stayed below the complete estimator fitted on half the molecules (0.510) and rebuilt on all of them (0.515), while keeping the programme’s amplitude above the raw counts. The per-cell comparator pools molecules within each inferred cell: nuclei segmented on the post-Xenium H&E with 2-*µ*m squares assigned by nearest-nucleus expansion (bin2cell[8]; Space Ranger 4.0 also uses nearest-nucleus expansion[16]) covered 82–94% of window bins, but a cell held about six training molecules on the 289-gene panel, so per-cell composition placed the programme worse than the Gaussian (0.209) and only a cell-typing step that pools cells of one cluster brought the best of twenty settings to FICTURE’s level (0.463 on all molecules), below the complete estimator in 12 of 12 units on every rank endpoint, also when the fields were rescored on the covered bins alone (Methods). The 5,000-gene experiment changes the allocation, not the estimator (Fig. 3c). The programme now had 0.23 training molecules per bin, and spatial borrowing alone could rank bins (edge Spearman 0.297, AUC 0.703) but not scale them: validation chose strong shrinkage and the fields kept 0.19 of the raw contrast. Molecular borrowing is what this regime needs, and it is what delivers: the factor components alone took the fields to 0.450, 0.810 and 0.70 of the raw contrast, 95% of the whole gain over spatial borrowing, with the correction and refit adding amplitude and held-out likelihood (10 and 12 of 12 units) rather than placement. Validation molecules put 0.83–1.00 of the epithelial genes’ weight on factor-informed components, almost all of it on corrected estimates whose spatial part is the bin itself, so that the estimate is the factor prediction adjusted by the bin’s own few molecules (Fig. 3d). The factor component was chosen on a one-bin neighbourhood, finer than any single gene could support, because its bin loadings are estimated from all genes at once. The complete estimator reached 0.458, 0.816, overlap 0.737, boundary distance 1.12 bins and 0.76 of the raw contrast, above the spatial-only fields and every reference in 12 of 12 units on the rank endpoints and in 11–12 of 12 on overlap, against a Gaussian that could not rank at all (0.002, 0.507). This is above FICTURE and above segmentation at every count budget and setting, in 12 of 12 units on every rank endpoint (best FICTURE with twice the molecules 0.404, 0.764, 0.689; Extended Data Table 1). Rebuilt on every molecule, the fields gave 0.462, 0.816 and 0.739. FICTURE’s boundary distance (0.90 bins) remained smaller and it kept the programme’s amplitude above raw (1.1) where the data-chosen allocation kept three quarters; that residual amplitude is the price of the per-bin mixture, which buys the held-out likelihood the factor component alone does not reach. Positional endpoints alone are compared across the two chemistries, since capture sensitivity after Xenium is reduced in these experiments[15].

The result replicates on a second lung tissue and panel. Four cores from four donors of a lung tissue microarray measured by Xenium and then by Visium HD[17] were registered through the shared post-Xenium H&E (Fig. 3e and Extended Data Fig. 4), fitted whole-core in tiles and scored on the alveolar programme. The complete estimator placed the programme better than the Gaussian, the global merge and the spatial-only fields in 12 of 12 core × seed units and in 4 of 4 donors (edge Spearman 0.241 against 0.198; AUC 0.685 against 0.661; contrast 0.85 against 0.70), with the ordering holding within every donor and under a one-bin registration shift. Absolute agreement is lower because the alveolar compartment is a network of septa one to two bins wide; these four donors replicate the measurement, not a biological claim.

### 2.4 Xenium fields against protein channels on the same section

The same-section comparison tests RNA against RNA; protein channels on the same section test placement against a chemically independent reference. In a Xenium section of clear cell renal cell carcinoma with 27-plex protein imaging on the same section and instrument [18], transcript coordinates and the protein channels share one frame, so no registration was needed. Three antigen/programme pairs were prespecified from the panel: CD31 against an endothelial programme (PECAM1, VWF), and PanCK and E-cadherin against an epithelial programme. Compartments were thresholded from the smoothed channel by fixed rules (Otsu for the broad PanCK stain, median plus three robust standard deviations for the sparse vessel and E-cadherin stains), and four 640-*µ*m windows were chosen on the PanCK image only.

Against CD31, the complete model beat the separately tuned Gaussian in 12 of 12 window × seed units on every endpoint: edge-zone AUC 0.840 against 0.798, contrast 0.90 against 0.48 of the held-out contrast, band profile error 0.016 against 0.180, and the held-out scores (Fig. 4a,b). Both sources mattered for these two sparse genes: spatial borrowing with per-gene shrinkage on the spatial component restored part of their amplitude (contrast 0.52 to 0.74), and the molecular source with its per-gene correction then raised placement and amplitude together (AUC 0.821 to 0.840, contrast to 0.90; 10–12 of 12 units over the spatial-only model). The epithelial pairs were not informative in this tissue and are reported as such: PanCK separated only a quarter to a third of bins by a few intensity units (AUC 0.60–0.63 for every estimator; complete model 0.621 against 0.602, contrast 0.72 against 0.47, 12 of 12 units), and the E-cadherin compartment covered 0.3–2% of bins and did not coincide with the epithelial programme for any estimator, including raw counts.

**Fig. 4.**
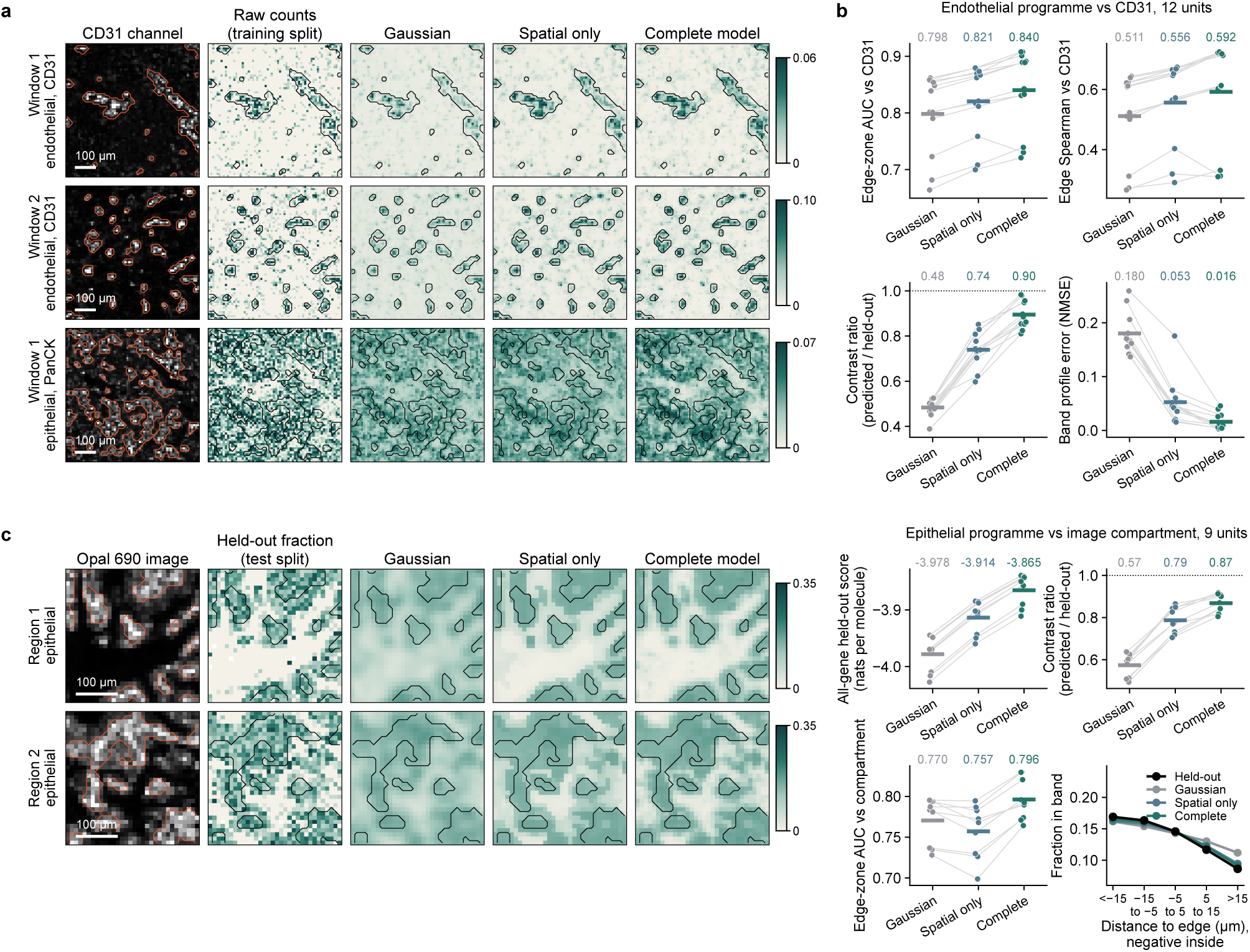
Xenium fields against protein channels on the same section. **a**, Clear cell renal cell carcinoma[18], two 640-*µ*m windows (10-*µ*m bins): CD31 channel with its thresholded image compartment outlined, raw training-split counts, the separately tuned Gaussian, the spatial-only model and the complete model for the endothelial programme (colour, programme fraction; contour, the same compartment); the PanCK pair is shown for comparison in the first window. Scale bars, 100 *µ*m. **b**, Endothelial programme against CD31, 12 window × seed units: edge-zone AUC, edge Spearman correlation with the channel, contrast ratio against held-out molecules and band profile error per estimator; grey lines join the same unit, bars mark unit means, numbers above give the means and the dotted line marks a contrast ratio of 1. **c**, Non-small-cell lung cancer section[19] with an Opal 690 image compartment (Otsu threshold), two regions of interest shown as 320-*µ*m zooms: image, held-out programme fraction from the test split, and the three fields for the epithelial programme. Right, 9 ROI × seed units: all-gene held-out score, contrast ratio and edge-zone AUC, showing the molecular gains and the rank deficit of spatial borrowing alone; and the pooled programme fraction by signed distance band from the compartment edge, held-out molecules against each estimator (unit means).

A lung cancer Xenium section with post-Xenium immunofluorescence[19] gave a narrower test against an unidentified dye channel whose positive compartment forms cords one to three bins wide; the complete model improved every endpoint over the Gaussian in 8–9 of 9 units (Fig. 4c), with the molecular source removing a deficit that spatial borrowing alone created in those thin cords.

### 2.5 A fibrosis cohort: the focus is different RNA, not more RNA, and its rim is a transcriptional transition

The tests above use one section each and ask whether a programme lands where an independent measurement finds it. A pulmonary fibrosis tissue microarray measured on Xenium with a 343-gene panel[17] asks the question at cohort scale and against pathology: 45 cores from 9 unaffected and 26 fibrosis donors, with 22 histological features drawn by a pathologist on registered H&E. Every polygon was mapped into the Xenium frame from the authors’ cell-to-annotation table alone (at least 0.99 of annotated cells inside their own polygon in 43 of 45 cores and 0.989 in a 44th), nine programmes and eight annotation/programme pairs were fixed before fitting, and each core was fitted whole in tiles on all molecules, assigned to a cell or not, with the same estimator, bank and selection protocol as every other dataset (Methods). Cores are the units and donor medians are reported alongside.

Against the polygons the complete estimator matched the held-out compartment contrast more closely than the tuned Gaussian (median 0.92 of the held-out contrast against 0.67, nearer to one in 149 of 173 core × pair units), with a smaller band-profile error in 152 of 173 and higher held-out scores in 172 and 173 of 173 (Fig. 5a,b). Rank agreement with the drawn boundary depended on the structure: the estimator ranked bins better for fibrosis against the fibrillar matrix programme (24 of 31 cores) and for honeycombing and remodelled epithelium against the basal programme (23 of 29), was close for fibroblastic foci (11 of 14), and for airways and alveoli the smoother Gaussian ranked bins as well or slightly better while losing every molecular endpoint. A polygon drawn at H&E resolution is a coarse reference for a 10-*µ*m field, and the sharper field pays for its calibration at the polygon’s edge; we report both outcomes.

**Fig. 5.**
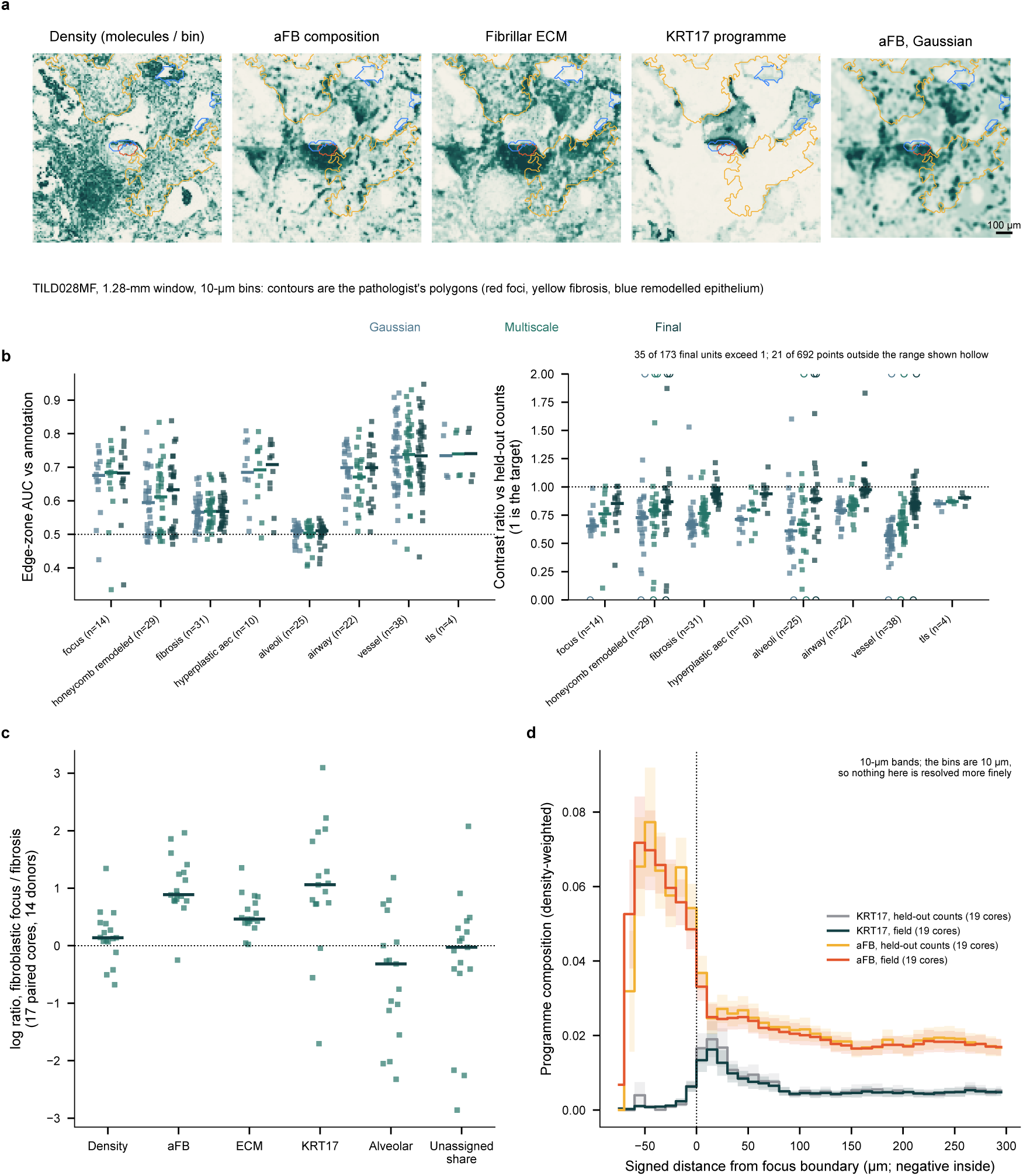
The fibroblastic focus is different RNA, not more RNA, and its rim is a transcriptional transition (45-core fibrosis cohort). **a**, One 1.28-mm window of a fibrosis core (TILD028MF) from the published cohort[17]: captured density (molecules per bin) and the activated fibroblast (aFB), fibrillar ECM and KRT17 programme compositions from the complete model, and the aFB composition from the separately tuned Gaussian. Contours are the pathologist’s polygons (red, fibroblastic focus; yellow, fibrosis; blue, remodelled epithelium). Colour scales are capped at the 99th percentile of each map. Scale bar 100 *µ*m. **b**, Edgezone AUC against the annotation boundary (left, dashed at 0.5) and contrast ratio of the compartment contrast to the held-out contrast (right, dashed at one; one is the target, not a ceiling) for the eight prespecified annotation/programme pairs; one point per core with seeds averaged, 173 core × pair units. Points exceeding the axis range (0–2) are drawn hollow on the axis; 35 of the 173 units for the complete model exceed one. Gaussian, spatial only and complete model. **c**, Log ratios of fibroblastic focus to surrounding fibrosis in the 17 cores (14 donors) containing both: fitted captured density, four programme compositions and the unassigned share of molecules; bars mark medians. The unassigned share is the only quantity that does not pass through a fitted field. **d**, Composition of the aFB and KRT17 programmes by signed distance from the focus boundary (10-*µ*m bands; negative inside, 19 cores with foci), from the fitted fields and from pooled held-out molecules; shading, s.e.m. across cores. The KRT17 rim rises just outside the boundary while the aFB programme falls away inside it, and the held-out counts track the field at every distance.

The fields also describe the focus itself, and here the comparison is one of calibration rather than of detection: pooled held-out molecules confirm the compartment-level direction independently, so the biological contrast does not depend on which estimator is chosen; what the per-bin field adds is the spatial structure that pooling cannot recover — the gradient across the boundary, the transitional bins, and the continuous map that supports spatial statistics. In the 17 cores (14 donors) that contain both a fibroblastic focus and annotated fibrosis, composition and density ranked the two compartments differently (Fig. 5c): the activated fibroblast programme (CTHRC1, POSTN, FAP)[17, 20] had 2.8-fold higher composition in foci than in the surrounding fibrosis (Hodges–Lehmann estimate, 17 cores, *P <* 0.0001; donor medians *P* = 0.0002) and the fibrillar matrix programme 1.7-fold (*P <* 0.0001), whereas captured density differed by only 1.15-fold and was not resolved at this number of donors. Segmented nuclear density provides a complementary measure of local packing. Computed from the Xenium nucleus segmentation on the same bins, it was 0.97-fold between the two compartments (17 cores from 14 donors, *P* = 0.28 on donor medians), as was the authors’ own cell assignment (0.91-fold). The two quantities are best compared as intervals, and they do not overlap: the compositional difference is at least 2.2-fold and nuclear density is at most 1.08-fold higher (Hodges–Lehmann estimates 2.79 and 0.92, 95% confidence intervals 2.24–3.75 and 0.73–1.08, 17 paired cores from 14 donors). The nuclear result is an upper bound set by small foci, which hold between 12 and 334 nuclei each, rather than a measurement of equal cell number. Captured molecules per segmented nucleus were 1.34-fold higher inside a focus (95% CI 1.16–1.53, *P* = 0.003). Within each core the captured-density ratio is exactly the product of the nuclear-density ratio and the ratio of captured molecules per nucleus, and the median of those per-core products is 1.15-fold, the same as the median captured-density ratio; the medians of the two factors taken separately summarise different cores and do not multiply. The observed contrast is therefore primarily compositional, with more captured RNA per segmented nucleus and no detected increase in segmented nuclear packing.

Two independent checks corroborate this. The compositional half does not need the estimator: assigning molecules to the authors’ own segmented cells and comparing compartments cell by cell gives the same answer (3.0-fold against the field’s 2.8-fold), and the small difference between them does not grow with the share of molecules belonging to no cell, taken over each core’s focus and fibrosis together and weighted by molecules, which spans 0.3 to 61.0% across these cores, so the field’s composition is corroborated by an orthogonal measurement rather than produced by it. The amount half is different: the same per-cell view puts the capture difference at 1.19-fold on donor medians and does not reject equality at that level (*P* = 0.08), and it cannot state a density at all, because its denominator is the segmentation. Pooled held-out counts gave the same ratios, so the field reproduces the compartment contrast without the polygon; across all 45 cores the fitted density tracked the segmented nuclear density at every scale (median Spearman correlation 0.62 at 10 *µ*m rising to 0.90 at 80 *µ*m), and no estimator differed from another there, so the fields neither invent nuclear structure nor smooth it away. The same comparison on a colorectal Visium HD section used an image-derived nuclear segmentation with no connection to the fitting: 305,411 of the 305,590 published H&E objects fell inside the RNA support, and their density per bin tracked the fitted captured density with Spearman correlations of 0.35, 0.52, 0.62, 0.69 and 0.80 across blocks of 8, 16, 40, 96 and 400 *µ*m, the fitted field, two Gaussian baselines and the raw counts agreeing to within 0.002 at every scale. A segmented nucleus is not a cell, so this constrains the field’s spatial pattern rather than its absolute rate. A focus is not a place with more captured RNA. It is a place with different RNA. The rim of the focus is a transition in transcriptional programme rather than in segmented nuclear density, with a profile measured in 20-*µ*m boundary bands (Fig. 5d). As a continuous profile that includes molecules assigned to no cell, the activated fibroblast programme fell from 0.051 inside the focus to 0.027 within 20 *µ*m outside it (half-decay distance below the 10-*µ*m band width), and the KRT17 programme formed a rim on the stromal side of the pathologist’s boundary, peaking at 10–20 *µ*m (0.0105 against 0.0029 inside and 0.0026 beyond 100 *µ*m; rim excess positive in 16 of 19 cores, *P* = 0.008). The authors described KRT5*^−^*/KRT17^+^ cells adjacent to activated fibroblasts from segmented cells[17]; the field resolves the same relationship as a 20-*µ*m band whose position and width are set by the molecules, and the held-out counts trace the same profile. All three estimators recover the rim and none of them creates it: its excess is 0.0032 in the held-out counts, 0.0029 in the fitted field and 0.0035 under the tuned Gaussian, each positive in 16 of 19 cores. The field’s band profile follows the held-out profile more closely than the Gaussian’s in 152 of 173 core × pair units; what the estimator adds is a continuous profile rather than detection. The unassigned share of molecules is a batch property of the two Xenium runs (0.2–5% in the first four arrays, 22–57% in the fifth) and is reported by batch, not interpreted.

### The same separation in a developing tissue

Density and composition were separated in the cohort by measuring both. A second specimen shows why the separation is not a formality: there the two quantities point in opposite directions. In an embryonic day 67 distal femur specimen [21] fitted whole with the same estimator at 16-*µ*m bins (88,814 bins, 25 tiles; held-out composition score 0.047 nats per molecule above a Gaussian refit on the same molecules, better in 25 of 25 tiles), the cartilage matrix programme (COL2A1, ACAN, COL9A1) had a 25-fold higher probability in the lowest-density third of bins than in the highest (2.5% against 0.10%; observed counts give 27-fold), while its captured intensity differed by only 1.3-fold between the two groups (0.0082 against 0.0103 molecules per *µ*m^2^, against a 37-fold difference in density). The low-density bins lie over the pale, rounded structures of the cartilage anlage. The direction is programme-specific rather than an artefact of the density split: collagen VI genes moved the other way, 1.4-fold higher in the dense bins (Extended Data Fig. 5). A larger share of the captured RNA therefore did not mean more RNA per unit area, and reading captured amount as expression would invert the comparison.

## 3 Discussion

Kintsugi estimates captured RNA density and gene composition on every observed tissue location, and it differs from earlier segmentation-free estimation in one decision: where each gene’s composition is borrowed from is chosen by held-out molecules rather than fixed by the method. Making that decision well took three components, and the ablations show each to be necessary. Spatial neighbours defined by total counts cannot see a composition boundary, so regions had to be built from the genes and their edges moved to where composition changes. One allocation for all genes suppresses the source that fits a programme best, so the allocation had to be per gene. A sparse gene has nothing to borrow spatially at any scale, so a molecular source had to be available, and validation molecules used it when it helped and left it unused when it did not: it was decisive on both real panels and in a 300-gene simulation with genuine co-expression, and it took 0.04–0.13 of the programme’s weight in a 16-gene simulation with nothing to borrow. The same simulations mark the boundary of the rule. The factor field’s geometry is chosen once for all genes, so when a programme needs a sharper neighbourhood than the rest of the panel its edge is placed worse even as the held-out score improves; held-out likelihood and edge accuracy are not the same objective, and we report that rather than tune against it.

The evidence has a definite shape. Positional agreement with an independent modality improved over a tuned Gaussian in all but one unit of the same-section and protein comparisons, and the fields’ compartment contrast approached the held-out contrast where the Gaussian halved it. Segmentation, the fixed allocation that borrows within the cell only, placed the programme no better than a Gaussian at six molecules per cell and reached FICTURE’s level only after borrowing across cells of a cluster.

FICTURE, a fixed allocation of molecular borrowing, sits between the ablations: below the spatial-only fields on the dense panel, above them on the sparse one, and below the data-chosen allocation on the rank endpoints of both, whether it is given the same molecules or twice as many. What it retains is a tighter boundary on the sparse panel and a programme amplitude above the raw counts, where the data-chosen fields keep three quarters of it and buy held-out likelihood with the difference.

Where the references are biological, the fields agree with what the molecules themselves say, and separating the two quantities changes the reading. A focus differs from the fibrosis around it several-fold in activated-fibroblast composition and by less than ten per cent in segmented nuclear density, the two intervals not overlapping, and it holds about a third more captured RNA per segmented nucleus; described by captured amount alone it is barely distinguishable, and described by composition it is a different tissue. The fitted fields reproduce those contrasts more closely than a tuned Gaussian does, and pooled held-out counts show them too, so at compartment scale this cohort tests calibration rather than revealing structure that pooling cannot reach; the estimator’s advantage is at the scale of a bin, where the same-section comparison measures it. The same separation appears in developing cartilage, where a higher programme share coexisted with a lower captured intensity, and in registered CODEX sections, where captured density carried information beyond segmented-cell count (Extended Data Figs. 5 and 6; the CODEX analysis is Supplementary Note 5). Captured amount at a tissue location combines its cell packing with its molecular state, and those two contributions can move independently: a focus has different RNA at nearly the same nuclear density, while the sparsest cartilage bins carry the highest programme share at the lowest captured intensity. Describing either tissue by captured amount alone would obscure the compositional change in the first and invert it in the second. Embedding and clustering methods (Banksy[22], SpaGCN[23]) and reference-based annotation or deconvolution (TopACT[14], cell2location[13]) primarily return domains, reference cell labels or cell-type abundances. Here the gene-composition comparators used per-cell or cell-cluster pooling and fixed configurations of a factor model; these comparisons sample the space of adaptive spatial methods.

Separating the two quantities opens questions that captured amount alone cannot frame: whether a treatment reshapes a niche’s transcriptional programme or its cell packing, and at what spatial scale a developmental boundary is compositional rather than proliferative. Because the borrowing decision is set by the data rather than by the assay geometry, it adapts as panels grow and resolution improves without redesigning the estimator.

The study’s scope is set by its references and its sampling. Thinned molecules evaluate prediction within a section, and only the fibrosis cohort offers donors as replication units for a biological claim; every reference compartment here, whether a thresholded dye image or a pathologist’s polygon, is itself an estimate at a coarser resolution than the fields, so none of these comparisons speaks below the scale of a bin. Matched imaging, replicated sampling and perturbation remain the ways to turn a well-placed field into a biological claim. The rule itself asks only for counts of genes at locations with an area: it needs no reference atlas, no segmentation and no assumption that a location holds one cell, so it applies to any assay that reports molecules on a support, at whatever resolution that support has.

## 4 Methods

### Data sources and native-bin support

Public count matrices and their coordinate tables were analysed without new tissue acquisition. The whole-section analysis included human CRC specimens P1, P2 and P5[1] at 8-*µ*m resolution and fetal knee cartilage[21]; native-bin widths for each extension were recorded with the input coordinates. The lung tissue microarray measured by Visium HD after Xenium[17] was analysed core by core as the second same-section comparison (below); its four cores were assigned to donors by matching the post-Xenium H&E to the per-core Xenium-registered H&E of the cohort, not by RNA.

Count features were aligned to the coordinate table by barcode. Nonnegative integer counts were retained in sparse form; duplicate gene symbols were combined while retaining the feature-to-gene mapping. The declared in-tissue mask defined the analysis support, including bins with no observed molecules. Rows were placed in mask raster order and the complete gene order was retained for fitting, saved models and scoring. Physical areas were calculated from native-bin width and actual in-mask support, not from bounding rectangles. Whole cores and sections were fitted in overlapping tiles and stitched by nearest tile centre (below).

### Separate estimation of captured density and gene composition

Let *Y_ig_* denote the observed molecule count for gene *g* in native bin *i*, with *T_i_* = ^Σ^*_g_ Y_ig_* and physical bin area *a_i_*. Captured RNA density *ρ_i_* is expressed in molecules per *µ*m^2^ at unit molecular exposure, and conditional gene probabilities satisfy *_g_ π_ig_* = 1. Under the working observation model *Y_ig_ ∼* Poisson(*e a_i_ρ_i_π_ig_*), where *e* is the retained molecular fraction, the likelihood factors into a Poisson term for *T_i_* and a multinomial term for gene allocation conditional on *T_i_*. The two factors may therefore use different spatial pooling. Throughout, *ρ_i_π_ig_* denotes gene-specific captured intensity per unit area. Neither density nor intensity is an absolute transcription rate or RNA abundance per cell.

Throughout, a molecule is one captured unit of the assay: a unique molecular identifier on Visium HD and a decoded transcript on Xenium.

Observed counts were divided into training (A), validation (B) and test (C) counts by sequential binomial thinning[24] with fractions 0.5, 0.25 and 0.25. Training counts constructed every candidate geometry and fitted every component. Validation counts selected parameters, components and mixture weights, after which the selected components were rebuilt on the combined training and validation counts (below). Test counts were scored only, and no test-count summary entered any choice. The implementation offers a further rebuild on all three splits, for producing a final field once evaluation is complete; it must be given the test counts explicitly, records that it used them, and is refused by the scoring routine. No field reported here was built that way. Under the working Poisson model, the three splits are independent Poisson draws with proportional rates; under negative-binomial sampling they are positively correlated, but a simulation showed that the estimator ordering and calibration were unchanged for dispersion *ϕ ≥*3 (Supplementary Note 6 and Extended Data Fig. 2). All comparisons here are between estimators on the same molecules. All estimators were evaluated on the same declared native-bin support and complete gene universe, including zero-count bins and undetected genes. Technical thinnings were not treated as biological replicates.

### Candidate geometries from the genes themselves

Three families of pooling candidates were built from training counts on the complete support. Fixed grids used square cells of side *w ∈ {*1, 2, 4, 8, 16, 32} bins, intersected with the support and split into four-connected components. Gaussian candidates used weights exp(−‖*x_i−_x_j_* ‖^2^*/*2*σ*^2^) within 4*σ* bins, *σ ∈* 0.5, 1, 2, 4, 8, 16 bins, restricted to the support and not rescaled. A spatially constant global profile was always available. Adaptive candidates were connected partitions built from the full training count cube rather than from total counts. Starting from native bins, a greedy merge tree joined the adjacent pair of regions whose union lost the least joint Poisson profile score,

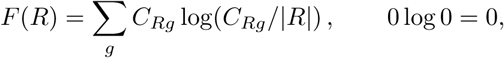

where *C_Rg_* is the training count of gene *g* in region *R* and *R* its area in bins; the pair minimising *F* (*R*) + *F* (*S*) *F* (*R S*) was merged, with equal costs ordered by region identifiers. Snapshots were retained at *K* 16, 32*, . . . ,* 4096 regions, restricted to *K ≤ N/*4 for *N* support bins. Because every gene contributes to the score, a boundary across which composition changes at constant density is visible to the merge; the same algorithm applied to total counts alone served as the control that isolates this contribution (Fig. 2).

Each snapshot was then refined by perimeter-penalised boundary moves. The objective was *_R_ F* (*R*) *− λ* per, where per is the number of four-neighbour bin pairs with different labels. Support bins were visited in four checkerboard classes so that moves within a class do not interact; a boundary bin moved to a four-adjacent region when the gain in *F* exceeded *λ* times the increase in perimeter, provided the donor region remained nonempty and four-connected (tested on the bin’s eight-neighbour ring). All accepted moves of a class were applied together and the class was reverted if the exact objective did not increase; sweeps continued until no move was accepted or 20 sweeps were reached. The penalty *λ* 3, 10, 30, 100 nats per boundary edge entered the candidate bank as four refined versions of every snapshot alongside the greedy snapshot, and validation counts chose among them exactly as they chose *K*. All geometry used training counts only.

### Component fitting and validation selection

For a candidate with nonnegative pooling weights *W_ij_*, define *S_i_* = Σ*_j_ W_ij_T_Aj_* and L_i_ = ∑j W_ij_ The density estimate was the Gamma-regularised posterior mean

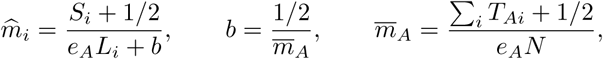

with *ρ̂_i_* = *m̂ _i_/a_i_*. Density candidates were compared by validation Poisson_Σ_log likeli-hood and one was selected. For composition, with global profile *q_g_*(*η*) = (∑*_i_ Y_Aig_* + *η/G*)*/*(∑*_i_ T_Ai_* + *η*), a candidate estimate was

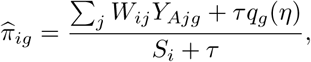

with *η* fixed at 0.5 throughout and *τ* 1, 10, 10^2^, 10^3^, 10^4^ selected jointly with the geometry by validation conditional multinomial log likelihood. Ties favoured the global endpoint, then regional before Gaussian candidates, then fewer regions or larger bandwidth, and then smaller *τ*. Within each family the best candidate was the family winner.

So that each gene can choose its own scale, each family also contributed a coarser sibling: the best-scoring candidate with at most *K/*4 regions (grid and adaptive families) or at least 4*σ* bandwidth (Gaussian family), or the coarsest candidate if none existed. Up to six spatial components therefore entered the mixture, before the molecular component below. Optionally, a per-gene shrinkage on a spatial component replaced that component’s single *τ* : each gene took its own *τ_g_* from 1, 3, 10*, . . . ,* 10^4^ , the component was renormalised within each bin, and the menu choice maximised the renormalised validation likelihood by a lagged minorisation–maximisation (MM) scheme[25] that bounds *T_i_* log *Z_i_*by its tangent at the previous normaliser *Z_i_*, which makes the choice separable across genes.

### The molecular source: factor components and factor-informed corrections

Molecular borrowing enters through two kinds of component. Training counts pooled into 4 × 4-bin units over the whole section or core (units with at least 20 molecules) were factorised once by Poisson non-negative matrix factorisation with multiplicative updates[26] (three seeded restarts screened at 40 iterations, the best continued to 200) into loadings *β_kg_*, rows normalised over genes, for rank *K* 8, 12, 20 ; every window or tile of that section shares these loadings, which are estimated from the whole support so that every window sees the same factor space. For every candidate geometry in the bank, not only the selected ones, the factor weights *θ_ik_* of every bin were the multinomial maximum-likelihood mixture of that geometry’s pooled training counts under the fixed loadings (30 multiplicative EM steps from uniform weights with a floor of 1*/K* per factor; regional geometries share one *θ* per region). The factor component 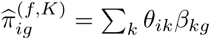 sums to one and needs no *τ* . Every (geometry, *K*) pair was scored by the validation conditional log likelihood; the best entered the mixture together with its coarser sibling under the rule above. Offering every geometry matters: on sparse panels validation chooses a finer neighbourhood for the factor component than for any single gene, because the factor weights are estimated from all genes at once. Sensitivity to the rank menu and unit size is reported in Supplementary Table 1. The winning factor field then served as the shrinkage centre of the spatial components. For each selected spatial geometry a factor-informed component replaced the global spectrum in the pooled estimator by the factor prediction for that bin,

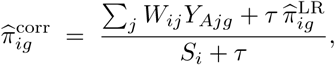

with *τ* chosen on validation counts from the same menu. The factor model states what a bin should contain; each gene’s own spatial neighbourhood corrects that by as much as its molecules justify. Because large *τ* drives the corrected component toward the factor field, the two factor-informed classes are near-duplicates and their sum is the determined quantity; across the three whole sections that sum carries 0.97–0.99 of the composition weight.

A multiplicative form of the correction, and a per-gene shrinkage inside the corrected components, which is a different option from the one on a spatial component above, were tested and are available in the package; their measurements are in Supplementary Note 8. The plain spatial components remain in the mixture: with small panels the factor prior is poor, and in the known-truth study removing them lost edge accuracy, so the mixture, not a fixed choice, decides between corrected and uncorrected estimates gene by gene.

### Refit after selection

Selection consumes the validation counts. Once every setting was fixed (density pooling, geometries, penalties, scales, *τ* , ranks, gene-wise shrinkage and weights), the selected components were rebuilt on the combined training and validation counts with those settings unchanged, and the per-bin normaliser recomputed; test counts were never used. Held-out scores and positional endpoints are reported for the refit fields unless stated otherwise, and the count-matched comparison with estimators that use all molecules rebuilds the components on all molecules for positional endpoints only.

### Gene-wise mixture weights

The components *π*^(*f*)^ were combined with a separate simplex weight vector per gene and renormalised within each bin,

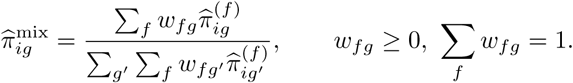

Global weights *u_f_* were first fitted by maximising the validation conditional log likelihood with sequential least-squares programming[27] after collapsing bit-identical fields, as before. Gene-wise weights then maximised the renormalised validation likelihood plus *α _f,g_ u_f_* log *w_fg_*, a pull toward the global weights with *α* = 1 molecule per gene. After renormalisation the objective is not concave; an MM scheme replaced *T_i_* log *Z_i_* by its tangent bound at the previous iterate, making the surrogate separable across genes with a closed-form fixed point per gene (one scalar root found by bisection). Fitting streamed over gene chunks for 200 iterations, the point at which the mixture amplitude had converged on the sparsest panel, with the returned normaliser recomputed for the returned weights. In the known-truth study *α* was instead chosen from 0, 1, 10, 100, 1000 by a half split of the validation counts; the choice shrank to global weights only in the smooth control. Programme probabilities are sums of gene probabilities over the fixed programme genes.

### Tiled fitting of whole cores

Cores and sections larger than about 2 10^4^ bins were fitted in overlapping square tiles (128 bins, step 96 bins; tiles with fewer than 400 support bins were dropped). Each tile was an independent unit of geometry construction, component fitting and validation selection on the shared thinning. Per-bin fields were stitched by assigning every bin to the tile whose centre was nearest, so that no bin was scored twice, and test-count scores were summed over owned bins. Tile boundaries can leave seams in the stitched field; all endpoints were computed on the stitched field without smoothing across seams.

### Computation

The estimator is implemented in the open-source Python package kintsugi (version 0.5.0). Pooled matrices of the selected geometries are cached once and reused across the mixture iterations, and the gene-wise weights are updated on a dense (gene, component, bin) tensor when memory allows, with a streaming fallback that gives the same result. On eight threads, one 16,384-bin window took 81 s with 300 genes, 756 s with 3,000 genes and 2,797 s with the complete 18,082-gene universe of a colorectal section; a 4,096-bin window took 14 s, 162 s and 1,635 s. Cost grows sublinearly in the number of genes (log–log slope 0.83) because the candidate geometries are shared across genes and the selected model has a fixed number of components. Fitting the section-wide factor loadings once took 58 s at 300 genes and 2,229 s at 18,082 genes on 545,913 bins. Peak memory is set by the dense weight tensor: below 6 GB up to 1,000 genes and 58 GB for 18,082 genes on a 16,384-bin window, which is why the budget is sized from available memory and the streaming path is kept. Tiles are independent and the driver’s stitching cost is negligible (0.02 s over nine tiles), so the honest unit of cost is the tile and a whole section costs the sum of its tiles; elapsed time depends only on how many tiles run at once. Three colorectal sections (507,684, 545,913 and 541,968 bins at 8 *µ*m; 72, 73 and 73 tiles; complete gene universe) summed to 49, 74 and 53 hours of tile fitting at 16 threads per tile, and the cartilage section (88,814 bins at 16 *µ*m, 25 tiles) to 20 hours; within a section the cost of a tile tracks its bin count, and over interior tiles of 12,000–17,000 bins the medians were 43, 69 and 52 min in the three colorectal sections (range 28–104 min). The cartilage section finished in 5.1 h of elapsed time with its 25 tiles over 80 CPUs; the colorectal sections finished in 5.2 to 7.3 h under a worker count that changed during the run, so those are wall times of particular runs rather than measurements at a fixed width, and the scaling of cost with tile size, gene count and worker count is measured separately. From a measured 47 min for a full 16,384-bin tile of the complete gene universe on one eight-thread worker, a 500,000-bin section is projected at 53 h on that worker and 5.2 h on twelve of them (Extended Data Fig. 7); the measured whole-section costs above are higher partly because their tiles ran twelve to a node, where the same tile took 1.7 times as long.

Tile seams were quantified by comparing, for each field, the median absolute difference between four-adjacent bins assigned to different tiles with that of interior pairs matched on distance from the tissue edge, with 95% intervals from a block bootstrap[28] over tiles (1,000 replicates) and a ratio of 1.0 indicating no seam; a pseudo-seam control repeated the measurement with the tile-ownership map translated by half a step, and the Xenium reference fraction, which never saw the tiling, served as a tile-free control. A seam was called only when the interval excluded 1 and the ratio exceeded that section’s own pseudo-seam control. Across the four lung cores the complete model’s steps were 1.1–4.1% of the field’s own range and did not exceed interior variation in three of the four, whereas the per-tile Gaussian exceeded it by 1.27– 2.79-fold; on the cartilage section the complete model showed no step (ratio 0.87, 95% CI 0.58–1.34; 0.10% of the field range) where the Gaussian exceeded interior variation 5.09-fold (95% CI 2.28–11.36). Log density showed no step for any estimator on the lung cores or the cartilage section (0.97–1.07, every interval containing 1). Where a step was called, it was a property of the tiling and not of the tissue: the excess was confined to the boundary line itself and returned to unity one bin away, and it was unchanged when interior pairs were additionally matched on local tissue roughness. A seam is therefore a boundary effect one bin wide, so the number of bins it touches scales with total boundary length; halving the tile edge roughly doubles it. Under the tiling used here, seam pairs are 0.8–1.2% of all adjacent pairs.

### Held-out and positional endpoints

Held-out scores retained the bin-by-gene count space: the conditional composition score of test counts per test molecule, computed over the whole support of a window or core, and, where a programme was prespecified, the binomial score of the programme’s test molecules against all others. Positional endpoints compared a programme probability field with an independent reference compartment that never entered fitting: Xenium molecules of the same section binned on the Visium HD grid, a protein channel of the same Xenium section, or a pathologist’s annotation polygons. A signed distance to the compartment boundary was computed on the support with half-bin correction, so that a bin touching the boundary sits half a bin from it, and with censoring of bins closer to the border of the field of view than to the boundary: the crop rectangle in the window and region-of-interest experiments, the tissue support itself in the whole-core experiments.

Within an edge zone (two bins, 16 *µ*m, against a Xenium reference; 15 *µ*m against a protein channel; 20 *µ*m against drawn polygons) we report three quantities. The area under the receiver operating curve (AUC) of the field against compartment membership, over every edge-zone bin including those where the reference detected nothing. The Spearman correlation of the field with the continuous reference, that is channel intensity, negative signed distance to a drawn boundary, or, against a Xenium reference, the Xenium programme fraction over the edge-zone bins in which Xenium detected at least one molecule; the same correlation over the whole window is also reported. And a binomial score of the field against the reference molecules themselves: the same-section Xenium counts where the reference is Xenium, the held-out test counts where the reference is a protein channel or a polygon.

Two further endpoints are taken over the whole support rather than the edge zone. The first is a contrast ratio, and two are used which are not interchangeable. Both are the difference in density-weighted programme composition between the two sides of the boundary, taken over bins near it, divided by the same difference for a reference set of counts. Against a Xenium reference, in the same-section and microarray experiments, the window is six bins (48 *µ*m) and the reference is the unpooled training counts, each bin carrying its own pseudocount-smoothed programme fraction, so raw counts score one by construction and a value below one is the amplitude the pooled estimator gives up; we call this the contrast relative to raw pooled counts. Against a protein channel or drawn polygons, in the renal carcinoma and fibrosis experiments, the window is 45 *µ*m and the reference is the held-out test counts pooled over the same bins, so one means the field reproduces the amplitude the held-out molecules show; we call this the held-out contrast ratio. The second is the normalised mean squared error of the field’s weight-pooled profile over signed-distance bands, five bands at 5 and 15 *µ*m against a protein channel and six at 5, 20 and 60 *µ*m against polygons, against the same profile of held-out test counts and normalised by the sum of squares of the reference profile.

In the two experiments with a Xenium reference, compartment overlap used an area-matched threshold: the field was thresholded at the quantile, over the bins surviving censoring, that reproduces the compartment’s area share among those bins, and intersection over union and the symmetric mean boundary distance in bins were computed on the resulting whole-support mask against the reference compartment.

A paired unit is a window, region of interest or core together with one thinning seed, the same split being given to every estimator compared; in the fibrosis cohort seeds are averaged within a core and the core is the unit. Comparators run on all molecules cannot be seed-matched and are scored once per window. A win is a strictly better value, ties counting for neither side: higher for rank, overlap and likelihood endpoints, lower for boundary distance and band error, and, for the cohort contrast, closer to one. Units are technical or spatial replicates, not independent specimens, unless stated.

### Known-truth simulation

Truth was a 32 × 32 lattice of 16 genes with a three-gene programme. Four scenarios set the programme probability to a smooth field (negative control), a step at constant density (composition-only), a step aligned with a four-fold density step, or a band five bins wide; nominal depth was 10 or 80 molecules per bin. Twenty replicates per scenario and depth drew independent Poisson training, validation and test counts at the same 0.5, 0.25 and 0.25 shares of the rate, which has the distribution binomial thinning would give. Geometry families were the previous total-count guide, a greedy Poisson merge of total counts, the joint merge of the full gene cube, and the two-category target merge; refinement penalties *λ* 0, 1, 3, 10 and merge sizes *K*doubling from 2 to 256 entered the bank. Endpoints were the mean absolute programme error in the two bins either side of the true boundary (edge error), the recovered contrast (the difference in mean fitted programme probability between the high- and low-programme regions of the field, divided by the same difference in the truth), the root mean squared error of the fitted probabilities of the thirteen non-programme genes, and the held-out programme binomial and all-gene scores, which in this experiment are lattice totals rather than per-molecule values. Comparators were the separately tuned Gaussian, a Gaussian tuned for the programme alone on validation counts, and an oracle that pools within the true regions. Paired wins count draws.

### Factor-structure known-truth simulations

A second known-truth study used a 64×64 lattice with every bin in the support and the boundary of the first scaled to it, *d*(*x*) = col (31.5 + 0.18(row 31.5) + 4 sin(row*/*12)), with the compartment *d >* 0 . Total density was flat at 10 or 80 molecules per bin, so every scenario is composition-only. Every gene entered every fit; no abundance filter was applied.

The 300-gene truth had eight latent programmes. Three scored genes carried the programme fraction in fixed shares 0.7, 0.2 and 0.1, and that fraction stepped from 0.015 to 0.090 across the boundary as in the sixteen-gene truth. Seven background programmes had Dirichlet(0.5) loadings over the 297 remaining genes and mixing weights *w*(*x*) on the seven-simplex; the eighth put a share *P_T_* of its loading on the three scored genes in the same split and 1 − *P_T_* on 30 further genes drawn without replacement with Dirichlet(1) loadings, which also carry background mass. The composition was *p*(*x*) = *θ*(*x*)*β_T_* + (1 *− θ*(*x*))*w*(*x*)*^⊤^β*_bg_ with *θ*(*x*) = *π*(*x*)*/P_T_* . Four scenarios: a flat background, *w* constant at one Dirichlet(3) draw with *P_T_* = 1*/*3, so 33 genes carry the step and the composition they share steps from 0.045 to 0.27; a structured background, *w* the softmax of that draw’s log weights plus seven standardised Gaussian random fields (white noise smoothed at *σ* = 8 bins), *P_T_* = 1*/*3; no co-members, the structured background with *P_T_* = 1, so the target programme loads on the three scored genes only and no other gene follows the step; and independent genes, each of the 297 background genes taking its own Dirichlet(0.5) base abundance times exp(0.7*z_g_*(*x*)) for its own standardised smooth field of the same length, renormalised to 1 − *π*(*x*), a full-rank truth with no co-members. The three scored genes step together in every scenario. One truth per scenario, from generator seed 2026.

The 1,000-gene truth had five co-expressed modules of 200 genes. Every gene drew a relative abundance log-uniformly over 0.1–50, 2.7 decades, and the loadings within a module were those abundances normalised, so realised gene depth ran from 8 10*^−^*^5^ to 0.084 molecules per bin at the lower depth and from 6 10*^−^*^4^ to 0.67 at the higher.

Module 0 stepped from 0.05 to 0.30 of the composition; the remaining share went to modules 1–4 through a softmax of one Dirichlet(3) draw and four smooth fields of the same length. A deviation variant departed from exact rank five in two ways, drawn once from the truth seed: half the genes of every module, 100 of 200 each, had their module field multiplied by exp(0.3*z_g_*(*x*)) for a standardised field of the same length, and a quarter of the step module’s genes, 50 of 200 drawn independently of the first set, had their own module share step to 0.05 + 0.25(1 ± 0.3) with a random sign, the composition being renormalised per bin afterwards. Generator seed 2027.

Counts were Poisson at the truth, drawn with seeds 0–19, twenty draws per truth and depth, and thinned once per draw into training, validation and test counts at 0.5, 0.25 and 0.25. Section-wide loadings were fitted by KL-NMF on the whole-lattice training split pooled into 4 × 4-bin units holding at least 20 molecules, all 256 units qualifying at both depths, at ranks 8, 12 and 20, three restarts screened at 40 iterations with the best continued to 200. The candidate bank was built from the training split alone: square grids of width 1, 2, 4, 8, 16 and 32 bins; a greedy joint Poisson merge of the full gene cube at *K* = 16, 32*, . . . ,* 1024, the menu 16–4096 restricted to *K ≤ n/*4; every snapshot refined at *λ* 3, 10, 30, 100 ; and Gaussian kernels *σ* 0.5, 1, 2, 4, 8, 16 bins. Forty-seven candidates entered every fit.

Four estimators were fitted, none tuned on truth. A Gaussian tuned for the programme alone took *σ* from that Gaussian menu plus a global pool and *τ* from 1, 10, 10 , 10 , 10 on the validation binomial score of the scored fraction, the module-0 total in the 1,000-gene truth, its density chosen separately on the validation Poisson score. The spatial-only stack was the published estimator with the factor family and the corrected components switched off. The complete estimator was the published preset: the factor family offered every scored geometry of the bank with the all-gene winner and its coarser sibling entering the mixture, one additive prior-shrunk correction per selected spatial component with *τ* chosen on validation counts, gene-wise weights, *η* = 0.5, the same five-value *τ* menu, and a rebuild of the selected components on the training and validation counts. A fourth adds one factor component of the winner’s rank on every selected spatial geometry not already carrying one. The 1,000-gene deviation variant omitted the fourth.

Endpoints on the scored fraction were the programme edge error, the mean absolute error over the bins with *d <* 2; the recovered contrast, the fitted high-minus-low difference over the same difference in the truth, taken over the two compartments entire; the root mean squared error over the low compartment; the boundary distance and the intersection over union of the field thresholded at the midpoint of the two levels, 0.0525 and 0.175 for the 1,000-gene module; and the held-out programme binomial and all-gene multinomial scores on the test split, here per test molecule rather than as lattice totals. Weight is the validation-selected gene-wise weight mass on the factor components, on the corrected components and on the remaining spatial components, averaged over the genes of a class. Per gene in the 1,000-gene truth, the relative edge error is the mean absolute error over the same edge bins divided by that gene’s own true step, and gene-depth deciles were cut on the distinct realised gene depths, over the 200 step-module genes for the per-gene endpoints and over all 1,000 genes for the weights, and are plotted at the geometric midpoint of each decile.

The 300-gene flat-background truth was also redrawn with negative-binomial counts of the same means at dispersion *ϕ ,* 10, 3, 1 independently per bin and gene, and with a Gamma(*ϕ, ϕ*) multiplier shared by all genes of a bin at *ϕ* 3, 1 , and passed through the unchanged pipeline, twenty draws per setting (Supplementary Note 6).

### Same-section Visium HD and Xenium

The 10x Genomics post-Xenium technical note describes two experiments on 5-*µ*m FFPE human lung adenocarcinoma sections from one donor. Within each experiment the same section was measured on Xenium (v1 289-gene human lung panel, Experiment 1; Xenium Prime 5K, Experiment 2) and then on Visium HD[15]. The Visium HD H&E and the Xenium-aligned H&E are the same scan: Space Ranger’s microns per pixel (0.2737) divided by the Xenium pixel (0.2125 *µ*m) equals the alignment-matrix scale (1.288). Xenium molecules (QV ≥ 20) were mapped through the vendor matrix onto the 8-*µ*m Visium HD grid; the direction of the transform was verified by total-count correlation on a coarse grid (matrix +0.54 and +0.49, inverse 0.23 and 0.51, in Experiments 1 and 2) and a whole-section integer-bin scan on smoothed log totals chose a ( 1, 1) bin offset. Shared genes with at least 500 Xenium molecules were retained (282 and 4,043); within each window the top 3,000 genes by Xenium count were kept with the programme genes forced in, so the fits used 282 and 3,000 genes per window. Four non-overlapping 128 × 128-bin windows were chosen by a rule on the Xenium epithelial fraction alone (25–75% epithelial bins at a single section-wide Otsu[29] threshold on the smoothed Xenium epithelial fraction, at least 95% tissue support, highest edge density). The epithelial programme was EPCAM, CDH1 and MUC1 (plus NKX2-1 in Experiment 2). The reference compartment in every window was that same section-wide threshold, with connected components smaller than four bins removed; the edge zone was two bins, 16 *µ*m. Seeds 101–103 gave 12 units per experiment. Estimators were raw training counts per bin, the separately tuned Gaussian, the joint merge with one global weight vector for all genes, the refined joint merge with gene-wise weights with and without multiscale stacking and per-gene shrinkage on a spatial component, and programme-versus-rest versions of the Gaussian and refined models, in which the programme’s genes are summed into one column and every remaining gene into a second, the estimator is run on that two-column matrix over the same training-count geometries, and each category’s fitted probability is divided equally among its members. The complete estimator adds the factor component on every selected geometry with an additive prior, section-wide loadings fitted once on the training split at ranks 8, 12 and 20, and a rebuild of the selected components on the training and validation counts; a count-matched variant rebuilds them on all molecules instead. FICTURE was run as punkst, its authors’ C++ implementation[12] (commit 8bd8ac5, 2026-04-20), on the whole section: hexagon widths 24 and 48 *µ*m, 8, 12 or 20 factors, three epochs, minimum hexagon count 20, pixel decoding at 8 *µ*m with anchor hexagons of 24 *µ*m and three moves, all factor probabilities retained. Two count versions were used: every shared molecule (FICTURE’s normal use) and the seed-101 training split (the molecules Kintsugi trains on). The composition at a bin is the decoded factor probabilities times the factor-by-gene loadings normalised over genes, and the programme probability its sum over the programme genes; contrast ratios weight bins by observed totals because FICTURE estimates no density. Bins without a decoded pixel (3–18% of window bins) were scored as zero programme probability. All six prespecified hexagon-width and factor-count settings are reported; the best on each endpoint is quoted as an upper bound within this menu. Kintsugi is reported with validation-selected settings only.

The segmentation-based comparator used bin2cell[8] (version 0.3.4) on the same sections: StarDist[30] nuclei on the full-resolution post-Xenium H&E that Space Ranger used, 2-*µ*m squares assigned to the nearest nucleus within an expansion distance (none, 4 *µ*m, the bin2cell default, or 8 *µ*m, the Space Ranger 4.0 default[16]), with bin2cell’s optional rescue of nuclei from the total-count image as a fourth label set. Space Ranger’s own segmented outputs exist only from version 4.0[31] and are not in the public download of this run (Space Ranger 3.0.0). Each cell’s programme fraction was computed from its assigned squares (pseudocount as for raw counts), on the seed- 101 to 103 training splits thinned at 2 *µ*m and on all molecules, and rasterised to the 8-*µ*m grid as the molecule-weighted mean over the cells in each bin; bins in no cell carried the global spectrum and are reported as uncovered. A cell-typing variant, added after the per-cell results were seen and labelled as such, clustered cells (Leiden[32] at three resolutions on the cell-level expression) and gave every cell its cluster’s pooled composition. Twenty settings were scored on the same windows, shift, compartment and endpoints as FICTURE, both with the setting chosen on validation molecules and with the setting chosen on the endpoint itself, and the Kintsugi fields were additionally rescored on the comparator’s covered bins alone. Coverage of window bins was 59–63% (nucleus only), 82–84% (4 *µ*m) and 92–94% (8 *µ*m).

### Xenium protein and image compartments

The Xenium Protein clear cell renal cell carcinoma dataset (405-gene panel, 27-plex protein, Xenium Onboard Analysis 4.0)[18] provides protein channels imaged on the same section and instrument as the transcripts, so bins were defined in one frame. Transcripts (QV ≥ 20, 62.9 M) were binned at 10 *µ*m; channels were read at the pyramid level nearest 2 *µ*m and averaged into the same bins. Four 64 × 64-bin windows were chosen on the PanCK image only (25–75% positive at the per-window Otsu threshold, at least 95% tissue coverage, a mean of at least 20 transcripts per bin, highest edge density, non-overlapping). Compartments used Otsu on the smoothed channel for PanCK and median plus three robust standard deviations for CD31 and E-cadherin, because Otsu returns an empty mask on near-unimodal channels; components smaller than four bins were removed. Programmes were fixed from panel membership (epithelial: EPCAM, KRT8, KRT18, KRT7, CDH1, MUC1, KRT5; endothelial: PECAM1, VWF). Seeds 101–103 gave 12 units.

The non-small-cell lung cancer Xenium section (patient L1, 289-gene lung panel) with post-Xenium immunofluorescence (DAPI, CD8, pan-cytokeratin) on the same slide[19] was analysed on three 64 × 64-bin regions of interest of 10 *µ*m chosen from the image before any RNA was examined, with the Opal 690 channel thresholded by Otsu (fixed threshold 40 as sensitivity) once per region on the unshifted channel, so that the nine registration shifts of up to ±10 *µ*m move the compartment without redefining it. The channel-to-antigen assignment is not stated in the public record, so the compartment is described as an image compartment; its extent (26–38% of bins as thin cords) is consistent with pan-cytokeratin rather than CD8, but this was not verified. The programme was EPCAM and KRT7.

### Lung tissue microarray measured twice

A post-Xenium Visium HD capture of one pulmonary fibrosis tissue microarray (GSE276934, sample GSM8509590, capture area IPFTMA5) covers four cores of the Xenium cohort above[17]: VUHD049 (unaffected), VUILD49 (chronic hypersensitivity pneumonitis), TILD111LF and TILD113MF (idiopathic pulmonary fibrosis), four 3-mm punches from four donors on tissue microarray 5. Each core was measured on Xenium with the 343-gene panel, stained, and then captured on Visium HD, so the two measurements are of one section. The two H&E images are the same post-Xenium stain: the whole-slide scan that Space Ranger aligned (0.5022 *µ*m per pixel, recovered from the positions fit) and the cohort’s per-core crops resampled into the Xenium pixel frame (0.2125 *µ*m per pixel). No landmark table, alignment matrix or cell assignment relates them, so the transform was fitted here from image content alone and RNA entered registration only through one integer-bin refinement.

The chain runs from a Visium HD 8-*µ*m bin centre to Xenium micrometres in three steps. First, bin centre to whole-slide pixel: the 2-*µ*m barcodes were aggregated in blocks of four (array row, array col integer-divided by 4) and the map from the 8-*µ*m array index to full-resolution pixels was fitted by least squares on the bins holding all sixteen members, which is an exact affine of the index (maximum residual 0.36 pixels). Second, whole-slide pixel to registered-core pixel: a similarity transform fitted from the images in two stages, a coarse stage matching the registered core at 8 *µ*m per pixel by normalised cross-correlation over the eight axis-aligned orientations, searched only inside the Visium HD capture area (whose position comes from tissue positions, not from counts), and a fine stage taking SIFT[33] keypoints at 2 *µ*m per pixel on the localised crop, ratio-test matches at 0.75, and a RANSAC[34] similarity with a 3-pixel (6-*µ*m) reprojection threshold. Third, registered-core pixel to Xenium micrometres: the cohort’s own translation-only registration of that image. Accuracy was measured by splitting the RANSAC inliers at random into halves, refitting on one and reporting the error on the other. Three cores registered this way (VUILD49, TILD113MF, TILD111LF): 22,292–24,836 inliers, inlier root mean squared error 1.74–1.89 *µ*m, held-out median landmark distance 1.26–1.36 *µ*m and 90th percentile 2.5–3.0 *µ*m, against an acceptance threshold of at least 100 inliers and a held-out median within one bin. The fourth core, VUHD049, has no registered H&E in the deposit; the three per-core transforms were each composed with their core’s translation into the Xenium slide frame, giving three estimates of one slide-level similarity that agreed to 0.04% in scale and 0.03*^◦^* in rotation, and their median was applied to VUHD049, displacing the slide centre by at most 2.3 *µ*m relative to any single estimate. That core is therefore registered by transfer and carries no landmark error of its own; it is also the only unaffected donor of the four.

Xenium transcripts (QV ≥ 20, gene features, whether assigned to a cell or not) were mapped back through the inverse chain and counted into the 8-*µ*m bins. The support is the core’s Xenium support carried onto the Visium HD grid, intersected with the closed, hole-filled set of bins holding at least one Visium HD molecule, reduced to its largest connected component (111,562–167,003 bins per core). Genes are the panel genes present in the Visium HD probe set with at least 500 Xenium molecules on that support, 252–280 per core. As in the first same-section experiment the registration was then refined by an integer-bin shift scan of ±3 bins on smoothed log total counts of all genes, totals only: ( 1, 1) for three cores and ( −1, 0) for TILD113MF. Total-count correlation at 8 *µ*m (0.49–0.60 on shared genes, 0.74–0.86 on smoothed all-gene totals) and the displacement against an RNA-only orientation-and-translation match were recorded for verification and never used to fit.

Each core was thinned once into training, validation and test counts (0.5/0.25/0.25) and fitted whole in 128-bin tiles at a step of 96 with nearest-centre stitching, one fit per core and seed, with the core-wide KL-NMF loadings (*K* 8, 12, 20 ) taken from that core’s training split. Estimators shared the tile’s training-only candidate bank, the validation split and the selection protocol, and were the raw training counts per bin with a pseudocount, the separately tuned Gaussian, the joint merge with one global weight vector for all genes, a spatial-only stack (refined merge, multiscale stacking, gene-wise weights, no factor component) and the complete estimator. The primary programme was alveolar epithelium (AGER, RTKN2, SFTPC, NAPSA, SFTPD), with KRT17 (KRT17, ITGB6, MMP7) and fibrillar ECM (COL1A1, COL1A2, COL3A1, DCN, LUM) as secondary compartment programmes; all were fixed from the cohort design before any fit. Each reference compartment was Otsu[29] on the smoothed Xenium fraction of its own programme, thresholded once per core independently of the thinning seed, with components smaller than four bins removed. Endpoints are those of the positional section, the contrast being the contrast relative to raw pooled counts. Seeds 101–103 on four cores gave 12 core × seed units; each core is a different donor, so the donor-level counts reported alongside are the four cores’ medians over seeds. Scores were also computed on 128-bin windows chosen within each core by the same rule as the first experiment (at least 95% support, 25–75% compartment bins, highest edge density, non-overlapping, quarter-window stride) as a secondary unit. Sensitivity to registration moved the chosen shift by one bin in both directions (+1, +1 and 1, 1) at seed 101 for the raw counts, the Gaussian, the spatial-only stack and the complete estimator.

### Pulmonary fibrosis cohort

Nuclear density was computed on the same 10-*µ*m bins from one centroid per segmented Xenium cell, taken as the mean position of that cell’s nucleus-overlapping transcripts. Every cell listed in the authors’ annotation table was recovered in all 45 cores and placed within 15 *µ*m of the position they report, and per-compartment counts agreed with their own cell-to-region assignment to within 17%. Captured density is not a cell count, so agreement with nuclei is a consistency check on spatial pattern rather than a validation of an absolute rate, and the Xenium nuclear segmentation is a proxy rather than an exhaustive census of biological nuclei.

GSE250346[17] provides 45 Xenium cores (343-gene panel) on five tissue microarrays from 9 unaffected and 26 fibrosis donors, ten of the cores being second or third cores of a donor: eight fibrosis donors contributed more than one core, five of them a more-affected and a less-affected core and three of them two cores of the same type, one of the eight contributing three cores in all, and one unaffected donor contributed two. The core identifiers are not a reliable guide to severity: three donors carry cores whose identifiers read less-affected and more-affected while the deposit’s sample type records two cores of one type, so severity was read from that column throughout, and pathologist annotations of 22 features as GeoJSON polygons in the registered H&E pixel frame together with a table assigning every annotated cell (Xenium centroid, annotation instance) to its region. The corrected gene names were verified from the data in one control core (ACTA2 six-fold enriched within 15 *µ*m of artery-annotated cells; MRC1 not). Transcripts (QV ≥ 20, gene features, assigned or not) were binned at 10 *µ*m and the core was identified from the agreement between listed cell centroids and the mean position of the transcripts assigned to each cell identifier (cell identifiers are not unique across cores). Polygons were mapped to the Xenium frame by one similarity transform per core: the Xenium pixel scale with translation from the median offset between polygon centroids and member-cell centroids, refined by coordinate search on the fraction of annotated cells falling inside their own transformed polygon (membership ≥ 0.99 in 43 of 45 cores and 0.989 in a 44th; the remaining core, at 0.90, used cell-derived masks). Nine programmes and eight annotation/programme pairs, the compartment definitions and the profile statistics were fixed before fitting and are listed in the design record. A core contributed a unit for a pair only with at least one instance and 200 bins in the ±20-*µ*m edge zone. Compartment comparisons (fibroblastic focus against fibrosis with foci excluded) used bins inside the polygons on the support; profiles used 10-*µ*m signed-distance bands from 100 to 300 *µ*m around each focus instance. The unassigned share of molecules differs by run (tissue microarrays 1–4 against 5) and is reported by batch. Cores were fitted with one thinning seed. On three cores refitted with two further seeds, edge-zone AUC and the all-gene held-out score changed by at most 0.02 and 0.005 nats per molecule between seeds for every estimator, whereas the contrast ratio varied by up to 0.3 for compartments with fewer than 500 bins; paired rankings between estimators were unchanged.

### Whole-section fits and comparisons

The three colorectal carcinoma sections (P1, P2, P5; 8-*µ*m bins, 507,684, 545,913 and 541,968 support bins, 18,082 genes) and the cartilage section were fitted with the tiled driver: one thinning of the whole section (seed 115000), section-wide loadings fitted once on the training counts, 128-bin tiles at a step of 96 bins with every bin owned by its nearest tile centre, and the same bank, selection, corrections and refit as in every window experiment except that the local-concentration menu included a sixth value (*τ* = 10^5^). The Gaussian comparator selected its composition scale and shrinkage on the same tiles by the same validation score and was refit on the combined training and validation counts. Held-out composition and joint scores were summed over owned bins. On the cartilage section (88,814 bins at 16 *µ*m, 25 tiles, 25.3 million held-out molecules) the complete estimator scored 3.189 nats per held-out molecule against

3.236 for the Gaussian refit on the same molecules and 3.245 for the Gaussian fitted on training counts alone, and it was better in 25 of 25 tiles on composition and on the joint score (gains of 0.125, 0.112 and 0.129 nats per molecule over the refit Gaussian on P1, P2 and P5, 209 of 209 tiles on composition and joint; Extended Data Fig. 8). Whole-section wall time was 5.2 to 7.3 h per section on 72 to 73 tiles in parallel (median 43 to 69 min per tile on 16 threads), and steps across tile boundaries were 1.8–2.5% of the field range for the complete model; the per-tile Gaussian’s absolute step was comparable but its seam-to-interior ratio was an order of magnitude larger because the smoother field has an order of magnitude less interior variation (Extended Data Fig. 9). Log density showed no step in two of the three sections; in the third, a step of 6% of the range was common to all estimators.

### Nuclear comparison on a colorectal section

Nuclei for P2 were the complete set of 305,590 objects in the published H&E segmentation archive[35], used as supplied with no expression-based filtering and including the 503 carrying no molecules. A projective map from array coordinates to full-resolution image pixels was fitted by least squares to the positions table and inverted to assign each centroid to an 8-*µ*m bin; it reproduced the supplied pixel coordinates to better than 10*^−^*^9^ pixels, and 305,411 centroids fell inside the 545,913-bin RNA support. Correlations are Spearman across square blocks of bins, over blocks lying entirely within the support.

### Singlet-selection accounting and the Figure 1 tissue window

Published P1, P2 and P5 annotations[1] defined the singlet-selection comparison independently of Kintsugi regions. The universe comprised in-tissue bins with non-missing Periphery annotations. RCTD[36] doublet-certain and doublet-uncertain calls were combined; absent and rejected calls were grouped as not assigned. The published deconvolution used a 100-UMI minimum, whereas the tissue universe retained lower-depth bins. Published UnsupervisedL1 labels defined tumour (Tumor), stroma (Fibroblast, Smooth Muscle, Endothelial) and immune (T cells, B cells, Myeloid) groups. Retention denominators were all bins in each group. Tissue-share denominators included all annotation groups, either before or after singlet selection.

Section P2 is fitted whole in 128-bin tiles at a step of 96 bins with every support bin owned by exactly one tile, so each interior tile owns a 96-by-96-bin square. Figure 1b displays the 9,216 bins owned by tile 15, rows 112–207 and columns 592–687 of the 838-by-838 grid, a 768-*µ*m square, and Fig. 1c describes the fit of that same tile, whose own extent is rows 96–223 and columns 576–703. The window is therefore the unit of the fit, and the two panels describe one model. Among the 40 fully covered tiles, this one was chosen before any fit as the tile whose observed epithelial fraction varies most between its fifth and ninety-fifth percentiles, which places normal mucosa and tumour in the same field. Panel b shows observed molecules per square micrometre, estimated captured density, the epithelial programme probability as a percentage of captured molecules, and their product as epithelial molecules per square micrometre; the programme is EPCAM, KRT8 and KRT19, KRT18 being absent from the 18,082-symbol universe. Observed and estimated density shared a display scale; the composition and intensity scales were labelled separately.

### Matrix-related RNA programmes

The cartilage section used fixed descriptive programmes: COL1A1, COL3A1, DCN and LUM (fibrillar matrix); EPCAM, KRT8 and KRT18 (epithelial); PTPRC, LST1 and CD3D (immune); COL2A1, ACAN and COL9A1 (cartilage matrix); and COL6A1, COL6A2 and COL6A3 (collagen VI). Programme membership and missing genes were retained explicitly: KRT18 is absent from this probe set, whose keratin series runs KRT1 and KRT10 to KRT17 and then KRT19, so the epithelial programme there is the sum over EPCAM and KRT8 and is reported as a two-gene field. This is a narrower list than the colorectal epithelial programme of EPCAM, KRT8 and KRT19 on the same gene universe: KRT19 is left out of the cartilage programme by the programme’s own definition, not by the platform. The lung microarray used its own programmes, defined from that study’s markers and listed in Supplementary Note 3, including a five-gene fibrillar-matrix programme that adds COL1A2 and is not the four-gene set above. Observed programme fractions and estimated probabilities were described separately, with fixed 30th/70th-percentile density groups and Spearman associations without bin-level significance tests. Observed fractions were undefined at zero-total bins, although those bins remained in the prediction support. Nuclear comparisons required verified bin-aligned measurements, which were unavailable for the cartilage section and the lung microarray. For E67 tissue context, the public GSM8993724 positions table matched all 175,561 rows of the analysis coordinate table, including all 88,814 supported bins. The supplied tissue image and image scale were sampled at registered bin centres without altering field values or the density groups. The original support, including bins over image background, was retained. No image-derived anatomical segmentation was introduced.

### Native-count comparison with CODEX

CODEX provides multiplexed protein imaging[37]. The supplied SPATCH COAD, HCC and OV RNA–CODEX serial-section pairs were analysed separately[38]. Integer, nonnegative RNA counts were verified throughout the sparse matrix. The supplied codex common RNA bins defined support, without filtering by expression-derived high-quality flags. Barcode coordinates were fitted to registered coordinates by affine least squares. Registered CODEX cell centres were mapped back to the native grid and retained only when their supplied common-support flag was true and their containing half-open bin lay in the RNA support. Affine residuals and physical step lengths were recorded; registration did not establish exact microscopic correspondence across serial sections.

Fixed 25 × 25 native-bin units had nominal 200-*µ*m sides. RNA density was the exact count sum divided by 64 *µ*m^2^ times the number of represented native bins. CODEX cell counts and marker means used the same bin union. Units with at least 313 supported bins and ten retained CODEX cells formed the common regression population. Segmented CODEX cells were a proxy for nuclear packing, not directly measured nuclei in the RNA section. Nested linear models included cell density alone, then epithelial-cell fraction, then the fixed 16-marker panel: CD8, CD20, CD3e, CD56, Pan-Cytokeratin, CD4, CD34, SMA, FOXP3, CD163, HLA-A, CD11c, MPO, CD68, HLA-DR and IDO1. Raw RNA density was primary and log-one-plus density was a specified scale sensitivity. Individual markers were also evaluated after cell-density and epithelial-fraction adjustment.

Fitted *R*^2^ and nested increments were complemented by out-of-fold *R*^2^ using spatially blocked validation[39]: five deterministic folds of 1-mm blocks. For block coordinates (*u, v*), the fold was (*u* + 2*v*) mod 5; no exclusion buffer separated neighbouring training and test blocks. Predictor centering and scaling were fitted within each training fold. Five hundred within-section block-bootstrap resamples[28] used seed 116000. These intervals described within-section variation, not patient-level uncertainty. Model increments were order-dependent associations; lineage-marker contributions were not interpreted uniquely as within-cell activity. No Kintsugi regions, region centroids or estimated fields supplied the RNA response in this comparison.

### Implementation and reproducibility

The analysis used Kintsugi 0.5.0. Sparse integer observations, complete support and gene order were validated at the estimator boundary. Gene-column caches were shared across pooling candidates; predictions were evaluated in gene chunks. Selected models were saved as sparse arrays, numeric arrays and JSON metadata and reloaded in a separate evaluation process. The package included checks against direct small-array likelihood calculations, count-conservation and connectivity tests, deterministic candidate and selection checks, and save–reload prediction equivalence. The public candidate, fit, save, load and score interfaces and package demonstrations were checked in a clean Python 3.14 environment with the declared dependencies.

## Data availability

The whole-section Visium HD datasets are colorectal carcinoma specimens P1, P2 and P5 from de Oliveira et al.[1], available through the 10x Genomics dataset portal (https://www.10xgenomics.com/datasets). Published CRC metadata from the same study supplied expression-inferred cell-type labels; these labels were not independent pathological annotations. The externally supplied H&E nuclear segmentation for P2 is from Kiessling et al.[35] (Zenodo, https://doi.org/10.5281/zenodo.11402686); its domain annotations combine image and marker information and were not used as an independent histological control.

The same-section lung adenocarcinoma data (Xenium with the 289-gene and Prime 5K panels, then Visium HD on the same section within each experiment) are the 10x Genomics post-Xenium technical-note datasets[15]. The clear cell renal cell carcinoma Xenium section with same-section protein imaging is the 10x Genomics Xenium Protein dataset[18]. The non-small-cell lung cancer Xenium section with post-Xenium immunofluorescence is from Bilous et al.[19]. The pulmonary fibrosis Xenium cohort (45 cores, 343-gene panel, pathologist annotations) is from Vannan et al.[17] (GEO GSE250346). The lung tissue microarray acquired with Visium HD after Xenium is from Vannan et al.[17] (GEO GSE276934, one capture area on microarray 5; four cores from four donors: one control, one chronic hypersensitivity pneumonitis, two IPF), with the same-section Xenium molecules and per-core registered H&E from GSE250346. Human fetal knee cartilage data (E67 distal femur, 16 *µ*m bins) are from Jachim et al.[21] (GEO GSE260926). Registered Visium HD and CODEX data for the COAD, HCC and ovarian SPATCH specimens are from Ren et al.[38] (https://spatch.pku-genomics.org). Input specifications accompanying the analysis scripts identify the source files, bin sizes and roles of these datasets.

## Code availability

Kintsugi is implemented in Python and distributed as kintsugi-st on PyPI. The analyses described here use version 0.5.0. Source code is available at https://github.com/cafferychen777/kintsugi.

## Acknowledgements

This work used the ACES advanced computing resource at Texas A&M High Performance Research Computing through allocation MTH260094 from the Advanced Cyberinfrastructure Coordination Ecosystem: Services & Support (ACCESS) programme. The authors also acknowledge the National Artificial Intelligence Research Resource Pilot and ACES through allocation NAIRR260389. Portions of this research were conducted with advanced computing resources provided by the Texas A&M Department of Statistics Arseven Computing Cluster and the Grace and FASTER clusters at Texas A&M High Performance Research Computing.

## Author contributions

C.Y. developed the method, performed the analyses, generated figures and wrote the initial manuscript. X.Z. and J.C. supervised the project, guided statistical and biological interpretation, and revised the manuscript. All authors approved the manuscript.

## Additional information

Correspondence and requests for materials should be addressed to Xianyang Zhang or Jun Chen.

