## Extended Data for "Kintsugi decides, gene by gene, where spatial transcriptomics borrows information"

**Extended Data Table 1 Every estimator and comparator on the same-section reference.** Mean over the four 1-mm windows of the same-section dataset[1] (12 window × seed units where three thinning seeds apply; comparators given all molecules are scored once per window) of the edge-zone Spearman correlation and AUC of the epithelial programme against the Xenium fraction, the area-matched overlap (IoU), the boundary distance in 8-$\mu$m bins and the programme contrast relative to raw pooled counts. Kintsugi rows use the training split (half of the molecules) except the count-matched row, which rebuilds the selected components on all molecules. Two Gaussian baselines are shown because what a comparator is allowed to know matters. Selecting the scale and shrinkage on the whole panel, as the estimator itself does, drives the Gaussian towards a near-constant field on the sparse panel. The programme-versus-rest rows instead sum the programme's genes into one column and every other gene into a second and fit and select on those two categories, which is what a careful user would do knowing the target in advance; the pair of them, one restricted to Gaussian kernels and one given the full geometry set, separates what knowing the target is worth from what gene-built regions are worth. FICTURE[2] (punkst, six settings of hexagon width and factor count) and the segmentation-based estimator (bin2cell[3], twenty settings of expansion distance, nucleus rescue and optional cell typing) are shown at their best setting chosen on the endpoint itself, an upper bound: highest for $\rho$, AUC and IoU, lowest for boundary distance, closest to one for contrast. The contrast column is normalised to the same raw pooled reference throughout. In the table, $\rho$ denotes Spearman correlation. Sources: research/lowrank_candidate_20260908, research/ficture_comparator_20260908, research/segmentation_comparator_20260909.

| | Experiment 1, 289 genes | | | | | Experiment 2, 5,000 genes | | | | |
|---|---|---|---|---|---|---|---|---|---|---|
| Estimator (molecules) | $\rho$ | AUC | IoU | Bound. | Contr. | $\rho$ | AUC | IoU | Bound. | Contr. |
| Raw pooled counts (A) | 0.086 | 0.559 | 0.429 | 1.58 | 1.00 | −0.150 | 0.411 | 0.308 | 1.46 | 1.00 |
| Gaussian, scale and shrinkage selected on all genes (A) | 0.252 | 0.666 | 0.564 | 1.58 | 0.53 | 0.002 | 0.507 | 0.391 | 1.45 | 0.02 |
| Programme versus rest, Gaussian kernels (A) | 0.374 | 0.768 | 0.751 | 1.51 | 0.85 | 0.177 | 0.636 | 0.587 | 2.38 | 0.49 |
| Joint merge, one global weight vector (A) | 0.279 | 0.685 | 0.599 | 1.54 | 0.51 | 0.149 | 0.605 | 0.517 | 1.35 | 0.03 |
| Programme versus rest, gene-built regions (A) | 0.428 | 0.800 | 0.777 | 1.42 | 0.88 | 0.277 | 0.699 | 0.632 | 2.21 | 0.57 |
| Spatial only (A) | 0.427 | 0.792 | 0.758 | 1.36 | 0.59 | 0.297 | 0.703 | 0.633 | 2.07 | 0.19 |
| + molecular source (A) | 0.508 | 0.842 | 0.798 | 1.05 | 0.86 | 0.450 | 0.810 | 0.733 | 1.13 | 0.70 |
| + correction (A) | 0.501 | 0.839 | 0.800 | 1.07 | 0.93 | 0.450 | 0.808 | 0.730 | 1.13 | 0.74 |
| Complete model, refit on A+B | 0.510 | 0.846 | 0.809 | 1.05 | 0.94 | 0.458 | 0.816 | 0.737 | 1.12 | 0.76 |
| Complete model, rebuilt on all molecules | 0.515 | 0.850 | 0.812 | 1.05 | 0.95 | 0.462 | 0.816 | 0.739 | 1.12 | 0.75 |
| FICTURE, best setting (A) | 0.432 | 0.769 | 0.721 | 1.19 | 1.18 | 0.341 | 0.718 | 0.611 | 1.01 | 1.00 |
| FICTURE, best setting (all) | 0.456 | 0.782 | 0.764 | 1.13 | 1.16 | 0.404 | 0.764 | 0.689 | 0.90 | 1.08 |
| Segmentation, per-cell composition, 4-$\mu$m expansion (A) | 0.209 | 0.635 | 0.558 | 1.58 | 0.90 | 0.109 | 0.571 | 0.474 | 1.43 | 0.93 |
| Segmentation, best of 20 settings (A) | 0.447 | 0.794 | 0.724 | 1.23 | 1.00 | 0.260 | 0.673 | 0.570 | 1.24 | 0.97 |
| Segmentation, best of 20 settings (all) | 0.463 | 0.806 | 0.743 | 1.17 | 0.97 | 0.367 | 0.740 | 0.644 | 1.10 | 0.98 |

### Extended Data Figures

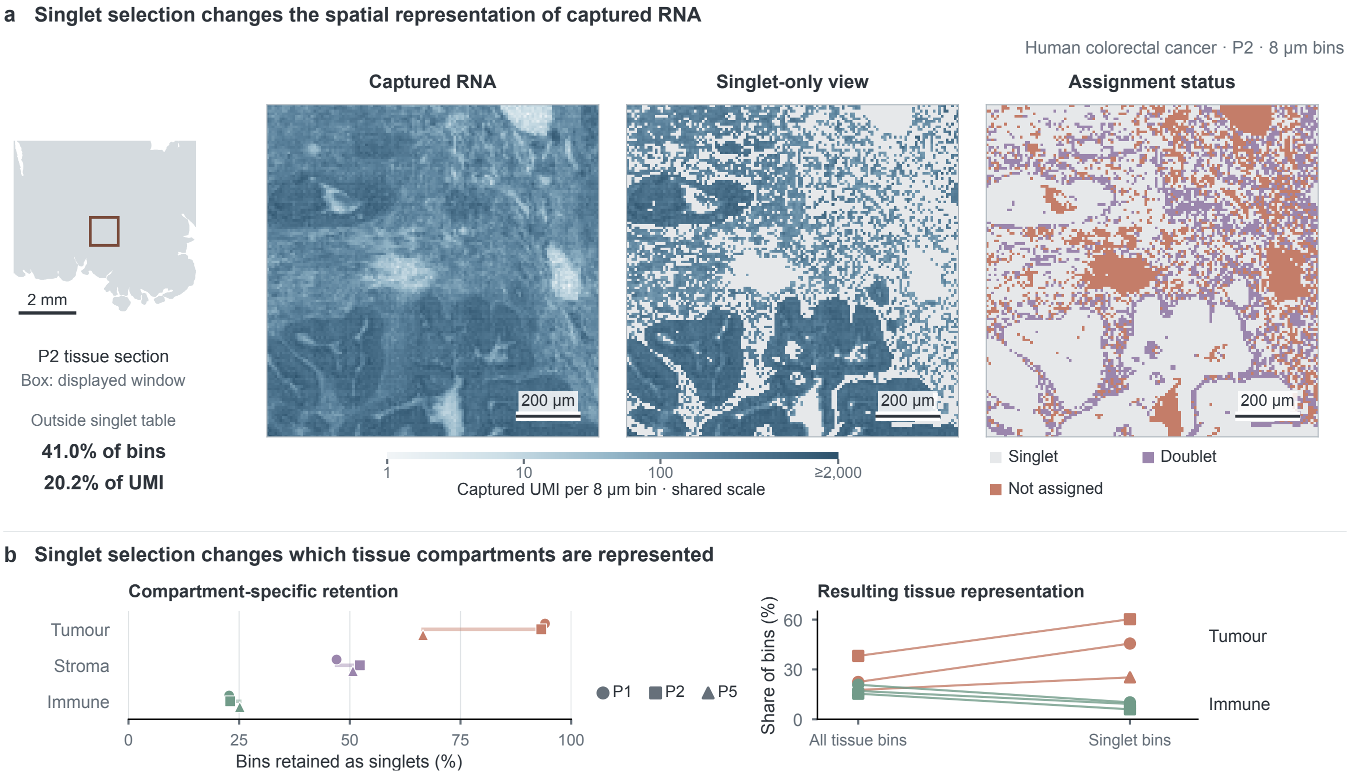


**Extended Data Fig. 1 | Singlet selection changes the representation of captured tissue RNA. a**, P2 colorectal cancer Visium HD data[4]. The locator identifies a 1.024-mm window; percentages describe the whole section. Observed UMI before and after published RCTD[5] singlet selection[4] share a logarithmic scale capped at 2,000 UMI per 8-$\mu$m bin. Grey indicates omitted bins. Assignment classes are deconvolution calls, not nuclear masks. Map scale bars, 200 $\mu$m; locator, 2 mm. **b**, Singlet retention within published tumour, stromal and immune annotations (left), and tumour/immune shares before and after selection (right). Symbols identify patients; paired points belong to the same patient. These are annotation-based bin fractions, not cell counts.

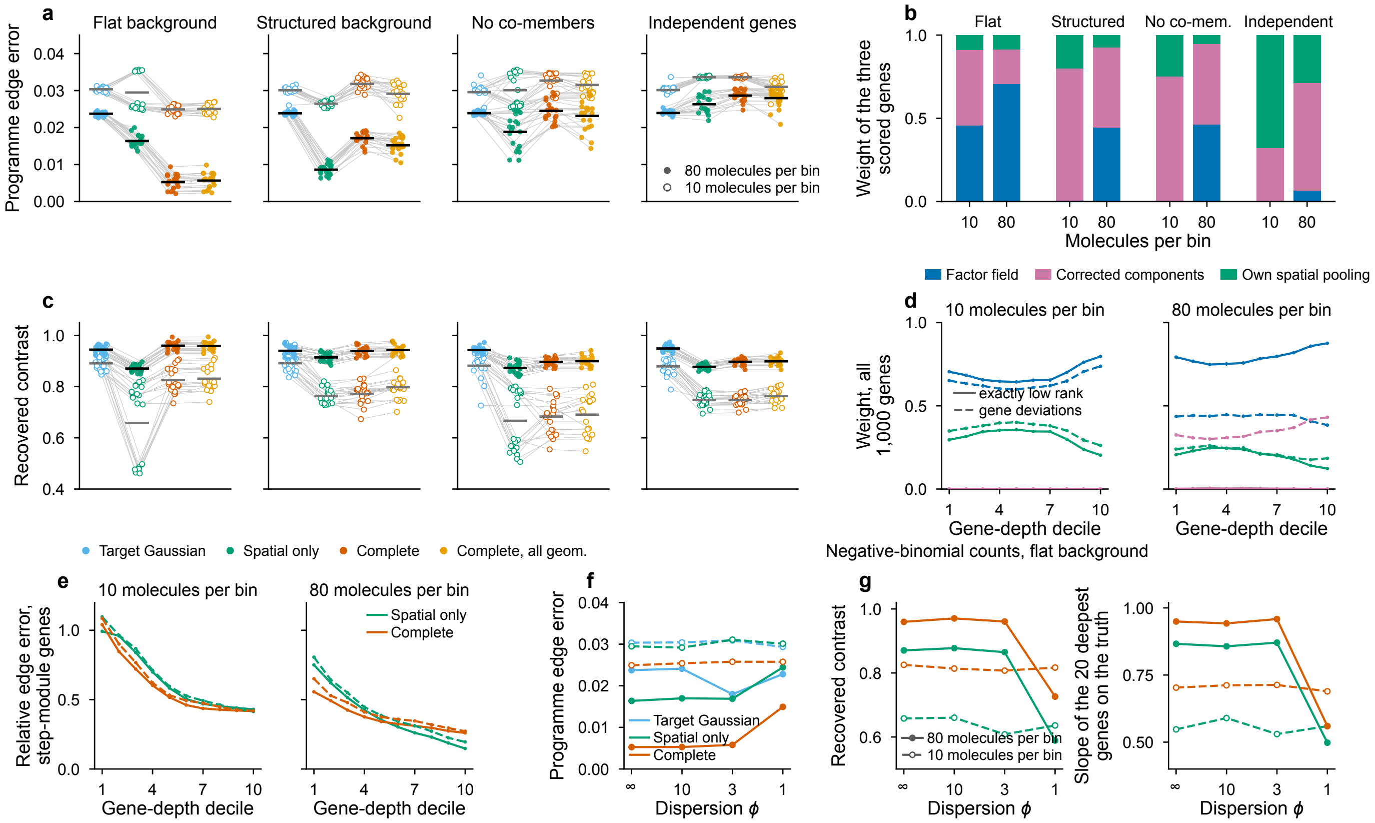


**Extended Data Fig. 2 | Known-truth simulations with genuine co-expression, and the two limits they expose. a**, **c**, Three hundred genes on a 64 × 64 lattice drawn from eight latent programmes with Dirichlet loadings. The three scored genes step together across a sinuous boundary at constant density in all four scenarios, which differ in the background and in whether the scored genes have co-expressed members. With a flat background and with a structured one, in which the seven other programmes vary smoothly across the field, the scored genes share their programme with 30 co-expressed members. In the third the target programme loads on the scored genes alone, so no other gene follows the step, over the same structured background. In the fourth the 297 background genes have independent rates and again no gene follows the step. Poisson counts at 10 and 80 molecules per bin, 20 draws each; thinning, candidate bank, loadings and selection as in the package. **a**, programme edge error, the mean absolute error of the three-gene fraction within two bins of the true boundary. **c**, recovered contrast, the fitted high-minus-low difference over the truth. Blue, a Gaussian tuned for the programme alone; green, the spatial-only stack; orange, the complete model; yellow, the complete model with the factor field offered on every selected geometry. Filled symbols, 80 molecules per bin; open symbols, 10; grey lines pair draws within a depth and horizontal bars are means. **b**, Mean validation-selected weight of the three scored genes on the factor field, on the corrected components and on their own spatial pooling, complete model, same draws. **d**, **e**, A 1,000-gene truth of five co-expressed modules of 200 genes whose relative abundances span 2.7 decades, so realised gene depth runs from $8 \times 10^{-5}$ to 0.08 molecules per bin at the lower

**Extended Data Fig. 2 | Continued**

depth and from $6 \times 10^{-4}$ to 0.67 at the higher one; one module steps across the boundary and the other four vary smoothly. Twenty draws per depth. Solid lines, a truth that is exactly rank 5; dashed lines, a variant in which half the genes carry a gene-specific smooth log-normal deviation from their module and a quarter of the step-module genes carry a step offset of ±30%. **d**, mean weight per source class over all 1,000 genes by decile of gene depth, complete model, colours as in **b**. **e**, mean relative edge error of the 200 step-module genes, each against its own true step. **f**, **g**, The 300-gene flat-background truth redrawn with negative-binomial counts of the same means and dispersion $\phi$ (variance $\mu + \mu^2/\phi$), independently per bin and gene, and passed through the unchanged pipeline; 20 draws per cell, and $\phi = \infty$ reproduces the Poisson harness exactly. **f**, programme edge error. **g**, recovered contrast and the slope of the 20 deepest genes' fitted proportions regressed on the truth across bins. The model's selection, ordering and calibration are unchanged for $\phi \geq 3$ and degrade only at $\phi = 1$ and 80 molecules per bin, where the deepest genes carry about one molecule per bin and the thinned splits are correlated at 0.24. The target-tuned Gaussian, which chooses its scale on the three scored genes alone, moves to a finer kernel already at $\phi = 3$ (blue, **f**).

The figure exists to show two limits. First, the factor family fits one geometry for all genes. With a structured background the all-gene validation chooses a smooth kernel; the scored genes still place 0.80–0.93 of their weight on factor-informed components, their all-gene held-out score improves in 20 of 20 draws at both depths and their contrast in 14 and 20 of 20, but their edge is placed worse than by spatial borrowing alone in 20 of 20 draws at both depths. Offering the factor field on every selected geometry reduces that loss without removing it: the excess over spatial borrowing falls from 0.0053 to 0.0027 at 10 molecules per bin, about half, and from 0.0085 to 0.0066 at 80, about a fifth, with 2 and 0 of 20 draws better than spatial borrowing. The same mechanism places the deepest step-module genes of the sparsity gradient worse than the spatial-only fields at 80 molecules per bin (**e**). Second, a truth that is exactly low rank leaves a gene nothing to gain from its own pooling at any depth, so weight on the factor field rises rather than falls with gene depth and the corrected components take no more than 0.005 of it (**d**, solid lines), the opposite of the allocation the same estimator makes on real panels. Gene-specific deviations switch the corrected components on at 80 molecules per bin, where they take 0.30–0.43 of the weight, but leave them at exactly zero at 10 molecules per bin.

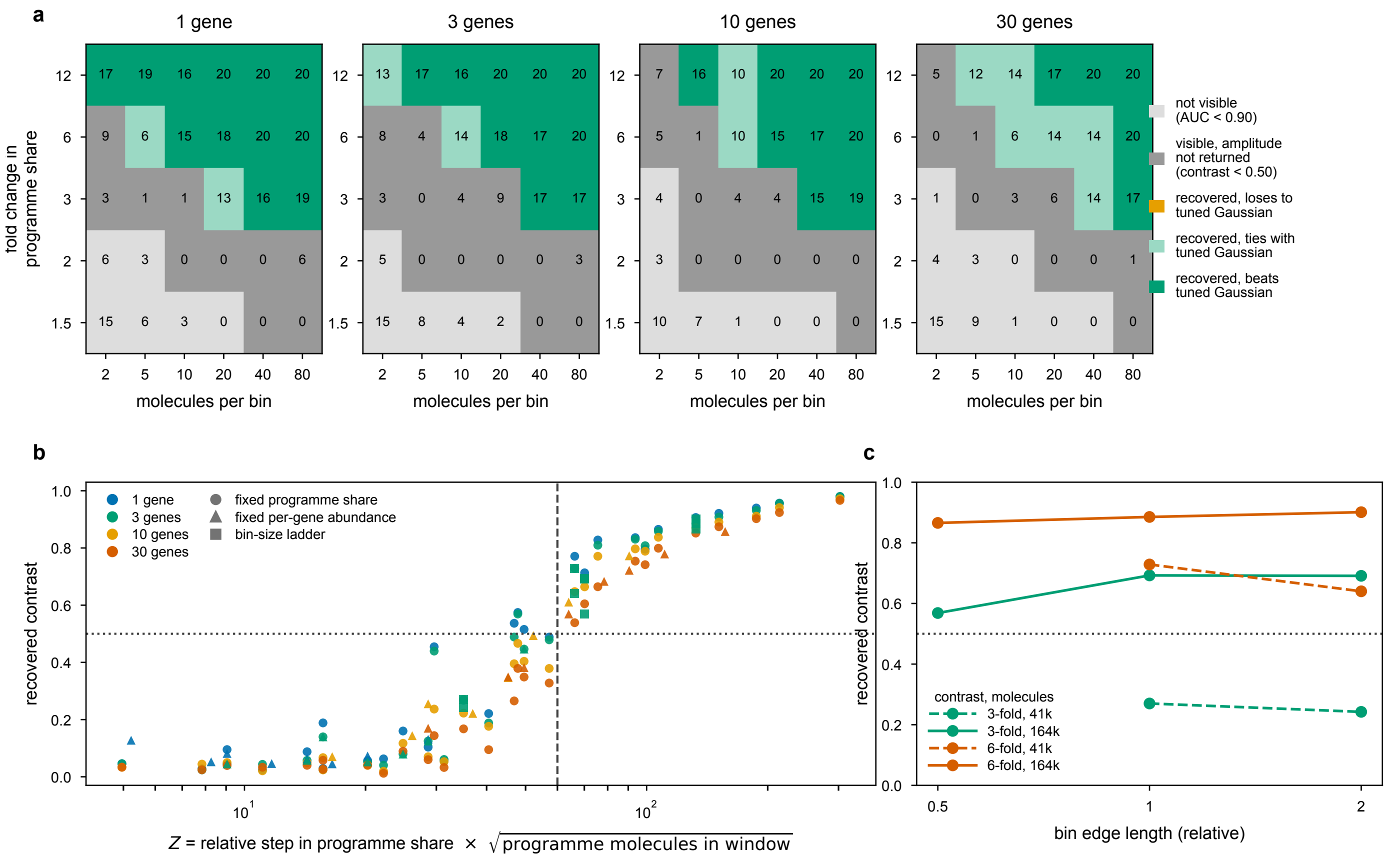


**Extended Data Fig. 3 | The estimator's operating range, and the statistic that organises it.** A 64×64 lattice of 300 genes drawn from eight latent programmes with a spatially flat background; a programme carried in equal shares by $M$ genes steps in composition across a sinuous boundary at constant total density (Methods). Twenty Poisson draws per setting. Estimators are the published preset, the same stack with the molecular source removed, and a Gaussian whose bandwidth and shrinkage are selected on that programme's own validation likelihood. **a**, Recovery verdict against sequencing depth and the fold change in programme share, one column per programme size $M$; the programme's share is fixed at 0.015 on the low side, so $M$ measures how many genes the same signal is divided among. Recovery is the prespecified criterion, mean AUC $\geq 0.90$ and mean recovered contrast $\geq 0.50$. Numbers in cells are paired draws out of 20 in which the estimator beat the tuned Gaussian on programme edge error, with $\geq 15$ counted as a win and $\leq 5$ as a loss. In every setting judged recoverable the estimator matched or beat the tuned Gaussian and no loss occurred (the orange class is unused); all 57 settings where it clearly fell behind lie outside the recoverable region, where neither method recovers the boundary. Panel **a** is one lattice geometry and its depth axis is a design axis rather than a general coordinate; the generalisation across bin sizes is carried by **b** and **c**. **b**, Recovered contrast against the window-level statistic $Z$, the relative step in programme share times the square root of the programme's molecules in the analysis window. Three datasets, 162 settings: the grid at fixed programme share (circles), a grid at fixed

**Extended Data Fig. 3 | Continued**

per-gene abundance in which the programme's molecule budget grows with $M$ (triangles), and the bin-size ladder of **c** (squares). Dashed line, $Z = 60$; dotted line, recovered contrast 0.50. The threshold misclassifies four settings, all of them conservative errors in which a boundary just below it was recovered anyway (recovered contrast 0.515–0.574). Fold change alone does not decide: a two-fold change from 0.015 to 0.030 was recovered at no depth and no programme size tested, while the same two-fold change from 0.150 to 0.300 was recovered from 10 molecules per bin, which is why the criterion has to be written in molecules rather than in folds. **c**, Bin-size ladder: one tissue, bin area varied, per-bin depth scaled inversely so that the molecules in the window are fixed, the feature keeping its physical size. Solid lines, 164,000 molecules; dashed, 41,000 (two bin edges only, because finer bins at that budget would hold fewer than three molecules). At a fixed molecule count a four-fold change in bin edge, and therefore a sixteen-fold change in per-bin depth, moves the recovered contrast from 0.57 to 0.69 at three-fold contrast and from 0.86 to 0.90 at six-fold, with no consistent direction; dropping the molecule count four-fold moves it to 0.24–0.27 and 0.64–0.73, which is the difference between recovering the boundary and not. **Domain of the rule.** Bin edge was varied over a four-fold range. The window holds one boundary spanning the field. Calibration is on flat backgrounds only; for how boundaries are placed when the background itself varies in space see Extended Data Fig. 2 and Supplementary Note 7.

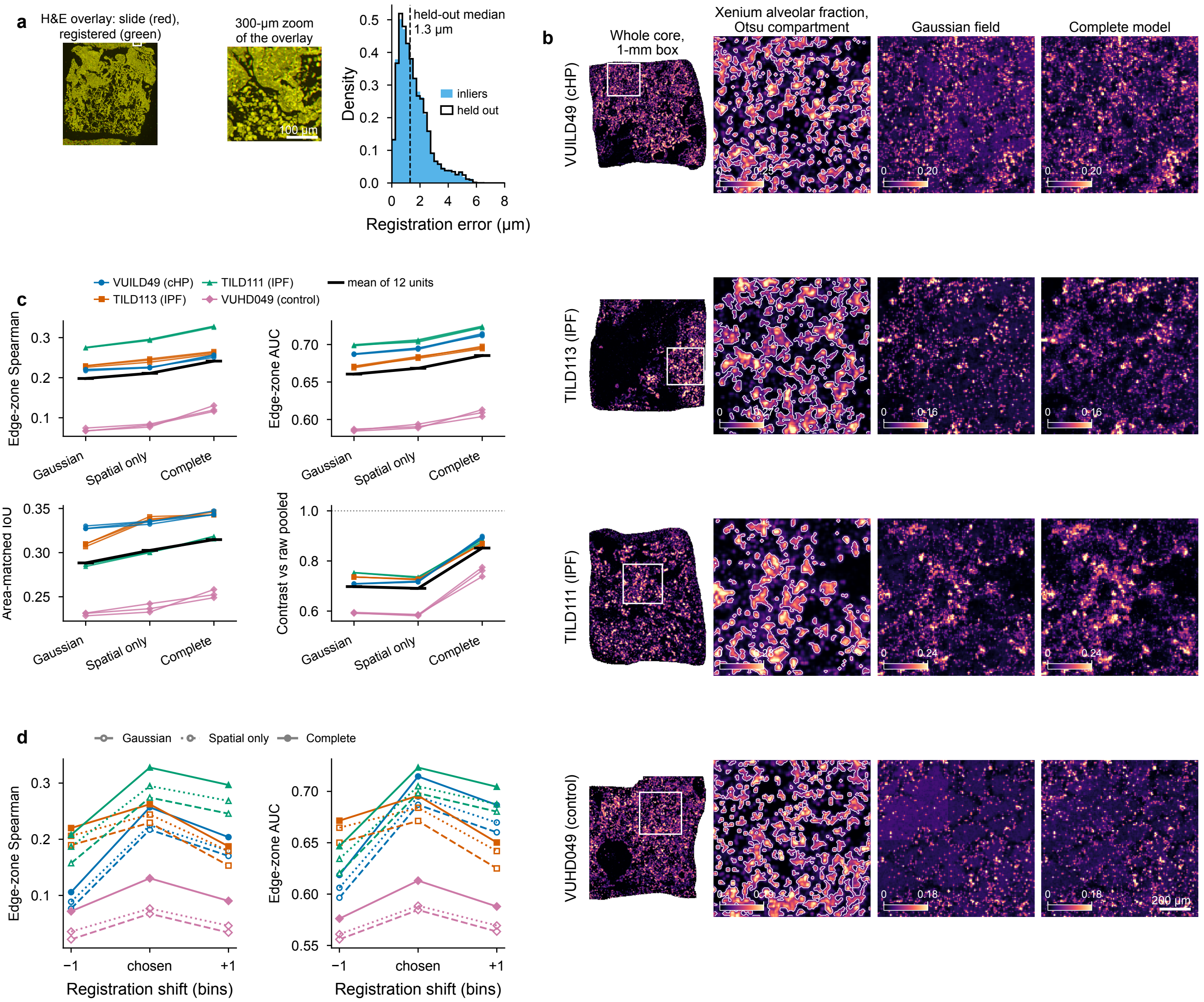


**Extended Data Fig. 4 | The same-section comparison replicates on four donors of a lung tissue microarray.** Four cores of one microarray section[6] measured by Xenium and then by Visium HD on the same tissue (one unaffected lung, one chronic hypersensitivity pneumonitis, two IPF). **a**, Registration from image content alone: the whole-slide post-Xenium H&E used by Space Ranger matched to the Xenium-registered H&E of the same core (template match, SIFT features[7], RANSAC similarity transform[8]), a 300-$\mu$m zoom of the overlay at the patch of highest H&E gradient, and the distribution of inlier residuals (filled) and held-out landmark errors (outline; median 1.3 $\mu$m). The three per-core slide-level transforms

**Extended Data Fig. 4 | Continued**

agree within 7 $\mu$m and their median was transferred to the core without a registered H&E; a one-bin shift chosen on total counts refines each core. **b**, One row per donor: whole-core Xenium alveolar-programme fraction on the Visium HD 8-$\mu$m grid with a 1-mm box placed by a rule (the window with the highest density of compartment boundary bins), then 1-mm zooms of the Xenium fraction (reference, Otsu compartment outlined in white, own colour scale), the separately tuned Gaussian field and the complete model (one shared scale per row), whole-core fits in tiles with core-wide loadings. Rectangular plateaus in the Gaussian zooms (clearest in the control core) are its per-tile scale selection stitched by nearest tile centre; the complete model shows a measurable step at tile boundaries in this core (seam-to-interior ratio 1.41 against the Gaussian's 2.15) but not in the other three (Extended Data Fig. 9). **c**, Paired endpoints over 12 core $\times$ seed units (edge-zone Spearman correlation and AUC against the Xenium fraction, area-matched overlap, contrast relative to raw pooled counts) for the Gaussian, the spatial-only fields and the complete model; lines join the same unit, symbol and colour mark the donor, black is the mean. **d**, Sensitivity to a one-bin registration shift in each direction (8 units, three estimators): the ordering Gaussian, spatial only, complete on the rank endpoints is unchanged. Units are technical replicates of four sections; the Xenium reference never enters fitting or selection.

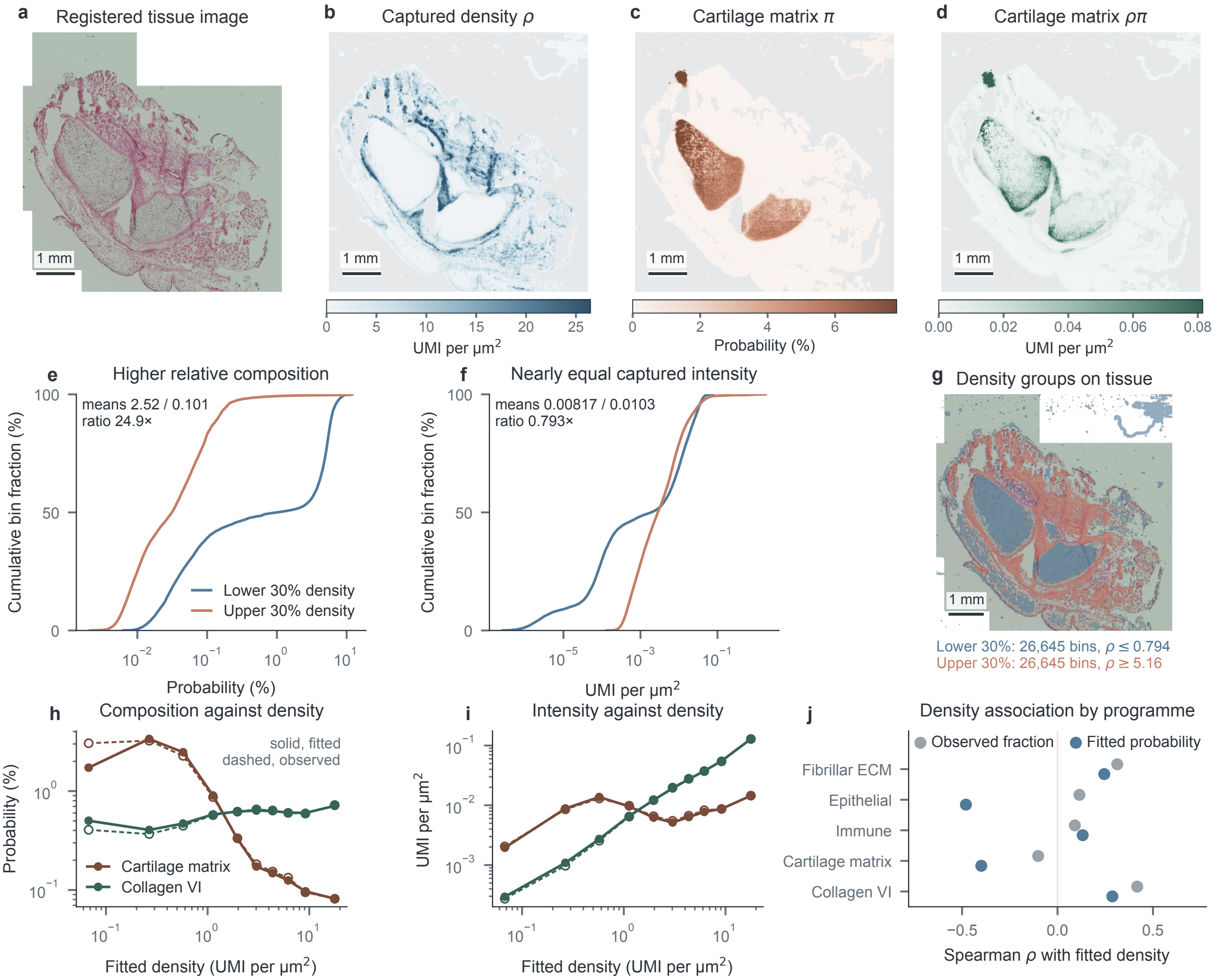


**Extended Data Fig. 5 | Relative composition and captured intensity diverge in developing cartilage.** Human embryonic day 67 distal femur (GSM8993724)[9], Visium HD, native 16-$\mu$m bins, fitted across the whole section with the tiled driver (25 tiles; all 88,814 supported bins retained). **a**, Registered tissue image at bin centres. **b–d**, Matched maps of fitted density $\rho$, cartilage matrix probability $\pi$ (COL2A1, ACAN, COL9A1) and captured intensity $\rho\pi$; each has its own labelled scale capped at its 99th percentile for display, and grey denotes outside support. **e,f**, Cumulative distributions of $\pi$ and of $\rho\pi$ in the lower and upper 30% of fitted density, with equal weight per bin and threshold ties retained (26,645 bins each; $\rho \leq 0.794$ and $\rho \geq 5.16$ UMI per $\mu m^2$). Annotated means and ratios use unclipped values. **g**, The same low-density (blue) and high-density (terracotta) groups on the registered image; these are defined by fitted density, not by anatomical annotation. **h,i**, Fitted (solid line,

**Extended Data Fig. 5 | Continued**

filled circles) and observed (dashed line, open circles) programme probability and captured intensity across deciles of fitted density, for the cartilage matrix programme and the collagen VI control (COL6A1, COL6A2, COL6A3); observed values are pooled counts within a decile. **j**, Spearman correlation of fitted density with the fitted probability (blue) and with the observed within-bin fraction (grey) for all five declared programmes; fitted values use all 88,814 bins, observed fractions the 87,809 bins with a nonzero total. Scale bars, 1 mm. Programme labels describe RNA identity, not extracellular origin. Curves and correlations describe within-section distributions, with no independent-bin tests or confidence intervals. The epithelial programme of panel **j** is the sum over EPCAM and KRT8 only: KRT18 is absent from this probe set and no alias of it is present, so a three-gene signature could not be measured here.

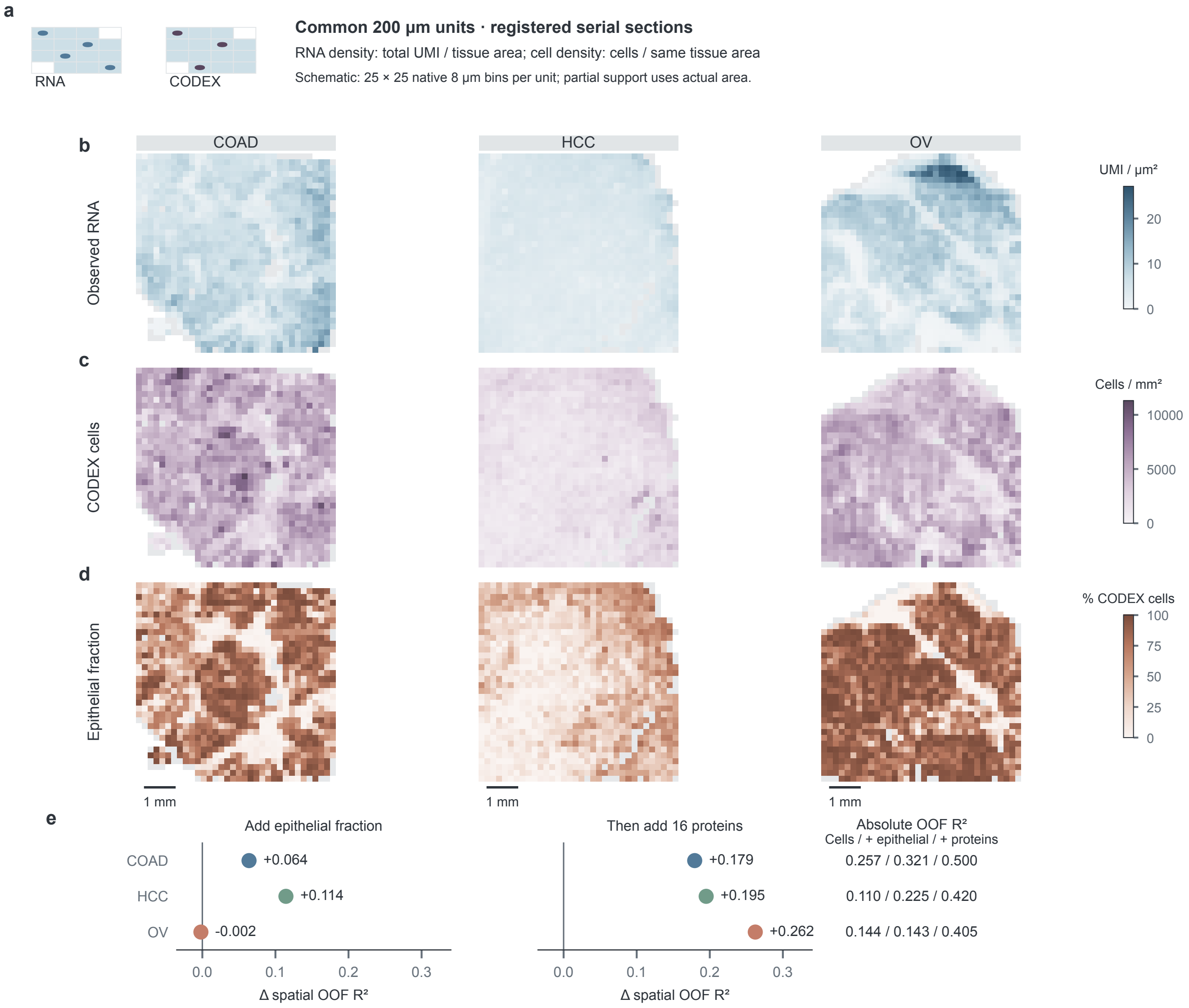


**Extended Data Fig. 6 | Cell density and protein composition jointly predict observed RNA density within serial sections.** **a**, Common nominal 200-$\mu$m units (schematic). Registered RNA and CODEX measurements share the same native 8-$\mu$m tissue-bin union. Density divides summed integer UMI or segmented cells by actual represented area, including partial units. **b**–**d**, SPATCH COAD, HCC and OV[10] (columns): observed RNA density (**b**), CODEX cell density (**c**), and the fraction of cells with published Epithelial annotations (**d**; not a protein image). Colours indicate regression-eligible units; grey, represented but excluded units; white, absent support. Unsmoothed maps share each row's full-range colour scale (fractions, 0–100%). Scale bars, 1 mm. **e**, Changes in spatial out-of-fold $R^2$ after adding epithelial fraction to cell density and then 16 prespecified proteins, in COAD, HCC

**Extended Data Fig. 6 | Continued**

and OV (1,063, 1,095 and 1,027 units). Columns share a zero reference and scale; adjacent text retains all three absolute scores. These are conditional prediction increments, not biological variance components. Five within-section folds use 1-mm blocks without an exclusion buffer. The epithelial-only addition slightly reduces OV performance. Segmented cells proxy nuclear abundance; serial sections do not measure identical cells. These specimen-specific observational comparisons are neither causal nor additive variance decompositions, and do not evaluate fitted Kintsugi fields. Supplementary tables retain fitted $R^2$ and bootstrap summaries.

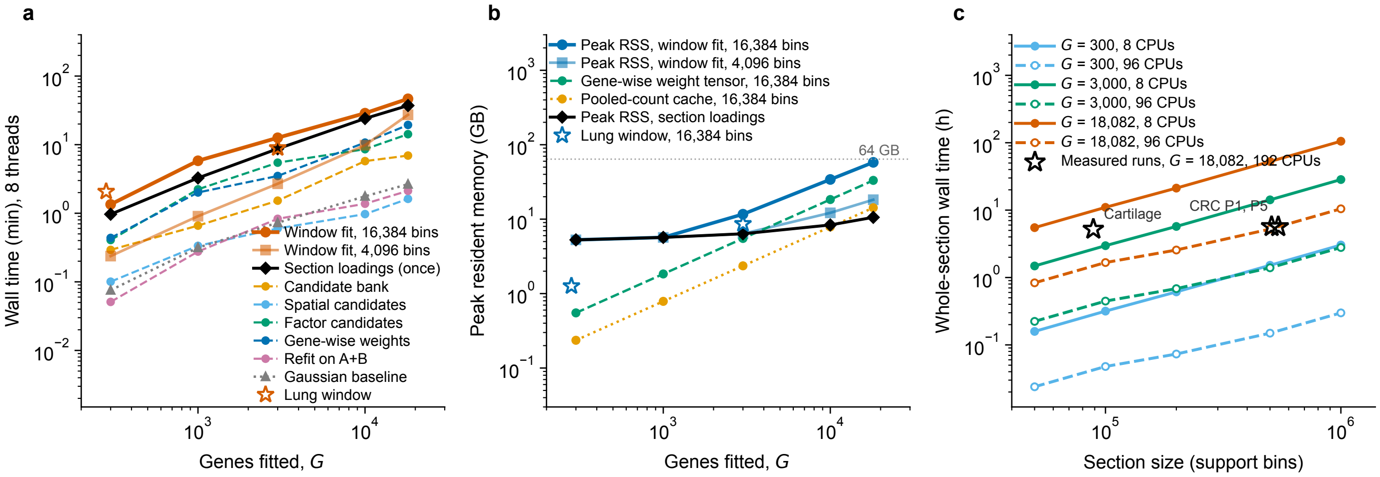


**Extended Data Fig. 7 | Runtime and memory of the estimator.** **a**, Wall time of each stage on eight threads against the number of genes fitted, measured on colorectal section P2 (Visium HD, 8-$\mu$m bins, 545,913 support bins, 18,082 genes). Solid orange, one 128 × 128-bin window (16,384 bins), comprising the candidate bank and the fit; pale orange, a 64 × 64-bin window; black, the section-wide factor loadings, fitted once per section over all bins and shared by every window; dashed, the five main stages of the 16,384-bin fit; dotted grey, an isotropic Gaussian composition baseline on the same window, without geometry selection. Stars, the two lung same-section slides (282 and 3,000 genes). Both axes logarithmic. **b**, Peak resident memory of the same runs. From about 3,000 genes the process peak is set by the dense gene-wise weight tensor (green) and the pooled-count cache (orange); below that it equals the resident section count matrix. Dotted grey, the 64-GB request used for whole sections. **c**, Whole-section wall time against section size. Lines, the projection loadings $+\lceil \text{tiles/workers} \rceil \times$ tile time, using the measured tile cost and the real tiling of P2 (73 tiles of 128 bins at step 96, 7,478 owned bins per tile); filled markers, one eight-thread worker on 8 CPUs; open markers, twelve concurrent eight-thread workers on 96 CPUs. Stars, three whole sections fitted with all 18,082 genes (colorectal P1 and P5 at 8 $\mu$m, fetal knee cartilage at 16 $\mu$m), run with 16 threads per tile across 192 and 80 CPUs; they fall above the 96-CPU projection because twelve tiles sharing a node contend for memory bandwidth, which cost the same tile a factor of 1.7, and because static shard assignment leaves stragglers (load-balanced ideal 4.6 h against 5.7 h measured for P1). The tiled driver itself is free: over nine tiles its wall time exceeded the sum of its tiles by 0.02 s.

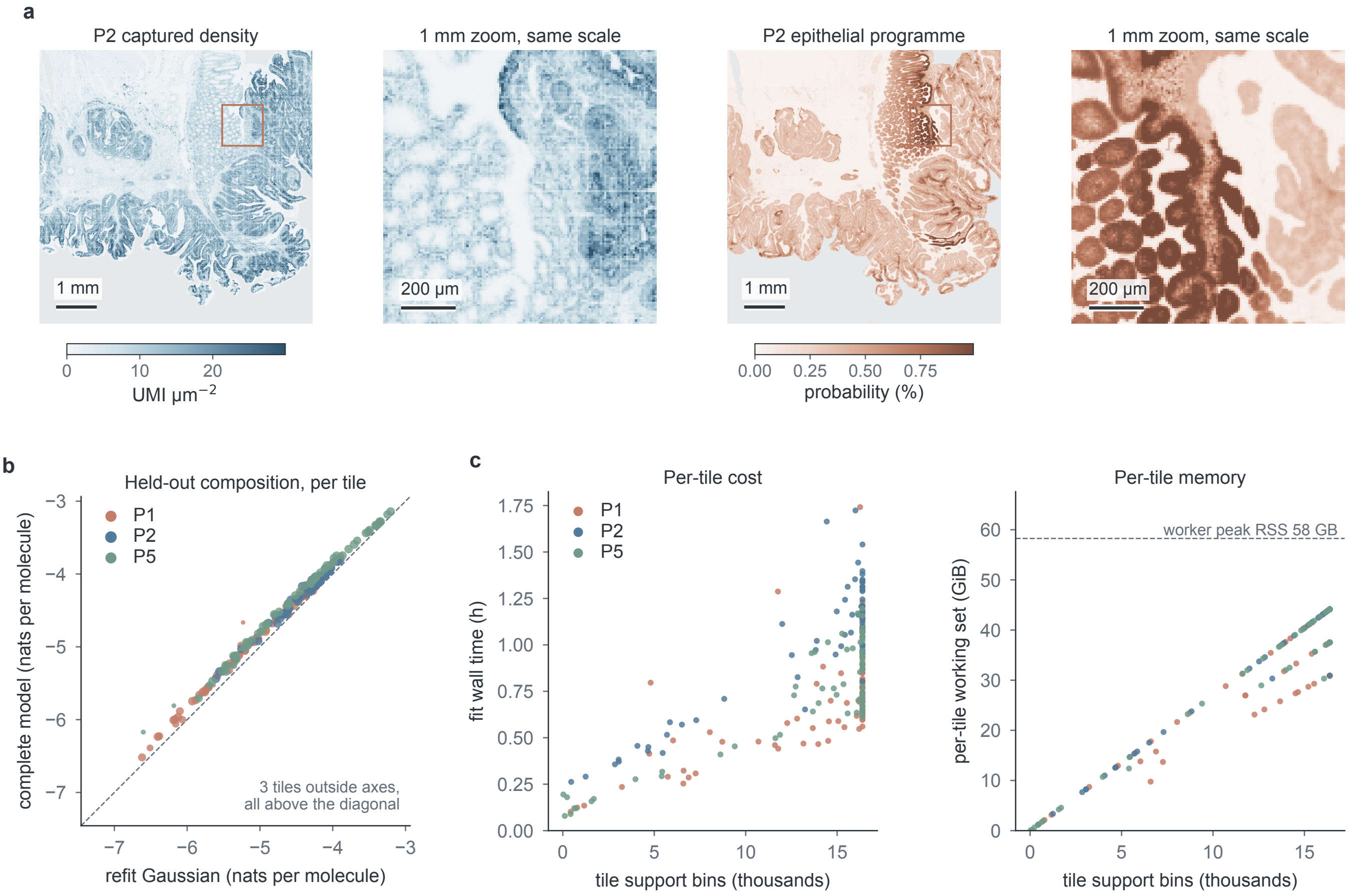


**Extended Data Fig. 8 | Whole-section colorectal fits replicate the held-out gains.** Three colorectal carcinoma Visium HD sections (P1, P2, P5; 8-$\mu$m bins, 507,684 to 545,913 support bins, 18,082 genes) fitted with the tiled driver: 72 to 73 tiles per section, section-wide loadings, and the same bank and refit as every window experiment. The complete estimator scored higher than the refit Gaussian on every evaluated tile (209 of 209 on composition and joint), with per-section gains of 0.125, 0.112 and 0.129 nats per held-out molecule.

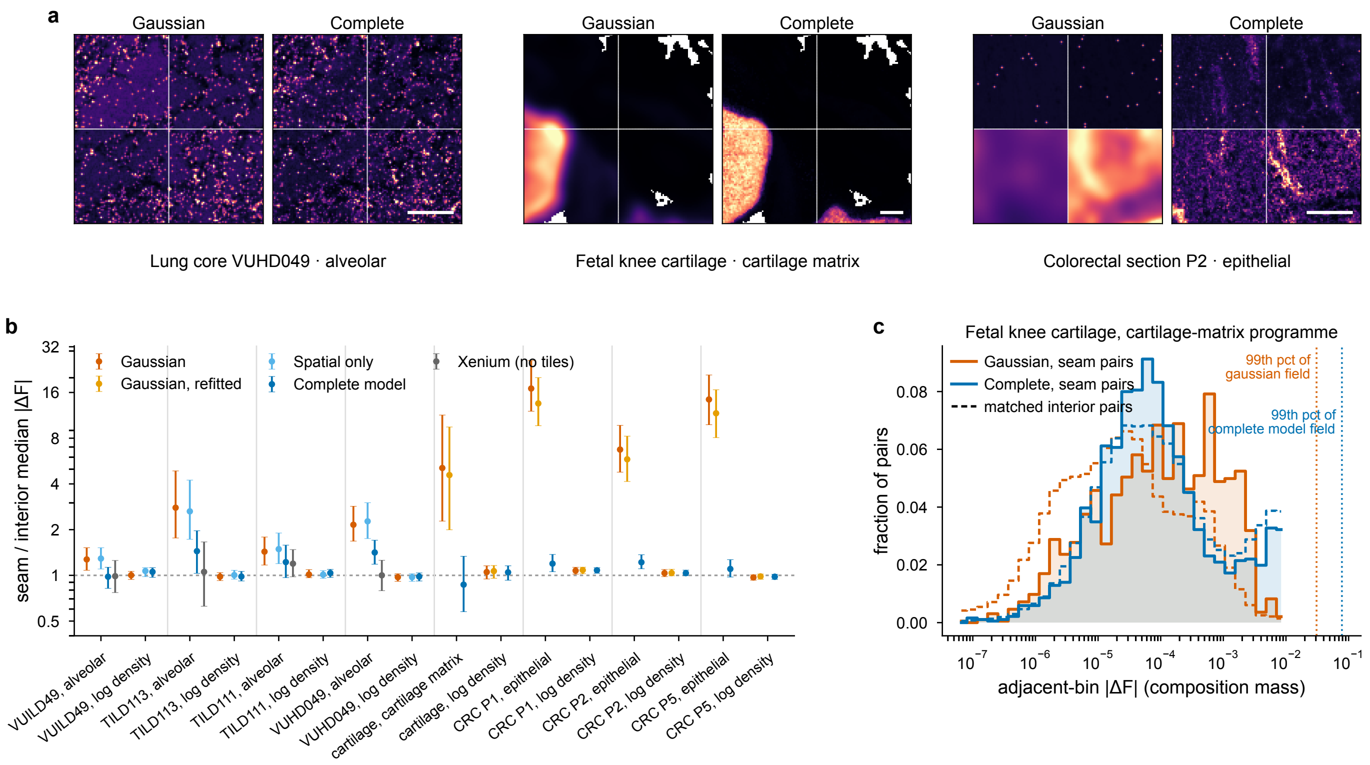


**Extended Data Fig. 9 | Discontinuity of stitched fields at tile boundaries.** Whole cores and whole sections are fitted in 128-bin tiles at a step of 96 bins and stitched by nearest tile centre. **a**, A 128-bin window placed by a fixed rule at a meeting point of tile ownership, for the control lung core (alveolar programme), the cartilage section (cartilage matrix programme) and colorectal section P2 (epithelial programme), showing the field of the Gaussian comparator and of the complete model; white lines are tile ownership boundaries, scale bar 250 $\mu$m. Each panel is scaled to its own 99.5th percentile, because the comparison is of steps within a field across a boundary, not of levels between estimators. **b**, Median absolute difference of adjacent bins that straddle a tile boundary, divided by that of interior pairs matched on distance from the tissue edge; one point per dataset and estimator, 1.0 meaning no seam, error bars 95% intervals from a block bootstrap over tiles (1,000 replicates). The Xenium fraction, which never saw this tiling, is a tile-free control, and a pseudo-seam control repeats the measurement on lines that are not tile boundaries. **c**, Distribution of absolute adjacent-bin differences for one dataset, solid for seam pairs and dashed for matched interior pairs, with the field's own 99th percentile marked; the Gaussian's seam pairs are shifted to the right and the complete model's two curves coincide. Seam pairs are 0.8–1.2% of all adjacent pairs. A large ratio is not a large artefact: the Gaussian's ratio of 5.1 on the cartilage section is a step of 0.36% of that field's range, while the complete model's ratio of 1.4 in the control lung core is a step of 4%.
