## Supplementary Information for "Kintsugi decides, gene by gene, where spatial transcriptomics borrows information"

#### Scope of the supplementary information

This supplement accompanies the Kintsugi 0.5.0 estimator described in Methods: geometries built from the training counts of all genes, refined by perimeter-penalised boundary moves; a molecular source in the form of factor components and factor-informed corrections; a separate simplex weight per gene chosen on validation molecules; and a rebuild of the selected components on the training and validation molecules once every choice is fixed. Note 1 states the identifiable targets and the score decomposition. Note 2 lists every fixed candidate menu and the evaluation units actually used. Note 3 defines the gene programmes and the observed-count controls. Note 4 summarises the alternatives that were tested and not adopted, together with the scope limits fixed by the known-truth study. Note 5 reports the native-count comparison with CODEX, Note 6 the robustness of thinning-based selection to overdispersion, Note 7 what the known-truth simulations do and do not reproduce, and Note 8 the alternatives to the corrected components that were implemented and left off by default, and Note 9 what the per-gene weights are and are not. Supplementary Table 1 reports the sensitivity of the factor menu, and Supplementary Tables 2–5 contain the native-count CODEX analysis, which does not depend on the estimator.

#### Note 1. Identifiable targets and score decomposition

For captured molecule counts, the working model is $Y_{ig} \sim \text{Poisson}(ea_i\rho_i\pi_{ig})$, where $a_i$ is native-bin area, $e$ is the retained molecular fraction of the split being scored, $\rho_i$ is captured molecules per unit area at unit molecular exposure, and $\sum_g \pi_{ig} = 1$. For $T_i = \sum_g Y_{ig}$, the density and conditional composition point scores are

$$\ell_{\rho,i} = T_i \log(ea_i\rho_i) - ea_i\rho_i - \log(T_i!),$$
$$\ell_{\pi,i} = \log(T_i!) - \sum_g \log(Y_{ig}!) + \sum_g Y_{ig} \log \pi_{ig}.$$

Their sum is exactly the original-bin gene Poisson point score, including all sample-space constants, because $\sum_g \pi_{ig} = 1$ and $\sum_g Y_{ig} = T_i$. The expected total per bin at unit exposure is $m_i = a_i \rho_i$, so that $\mathbb{E}[T_i] = e\, m_i$; the implemented estimate is $\widehat{m}_i$ and division by physical bin area converts it to density, $\widehat{\rho}_i = \widehat{m}_i / a_i$. Composition probabilities sum to one across the complete input gene universe, including genes unseen in training. Every shrinkage target is strictly positive (the global profile carries $\eta/G$ per gene and the factor prediction is a positive mixture of positive loadings), and the gene-wise mixture is renormalised within each bin, so no gene receives probability zero.

This factorisation does not make total counts sufficient for gene composition, justify ignoring composition boundaries, or establish that a data-selected partition retains all spatial information. Region sums are sufficient for a constant rate vector only conditional on that regional model. Different spatial regularisation can be appropriate for total captured amount and relative gene allocation, and the multinomial term constrains only the normalisation of $\pi_{i\cdot}$, so the composition estimator is free to borrow across genes at the same location (the factor components of Methods) as well as across locations.

If an unobserved capture efficiency $c_i$ multiplies biological RNA abundance per area $u_i$, the observation model identifies $\rho_i = c_i u_i$, not the two factors separately. Neither captured density nor an unadjusted correlation with it estimates absolute transcription, RNA per cell, or a bound on capture-efficiency effects. Protein and nuclear observations provide additional descriptive controls, not automatic identification of these latent factors.

### Note 2. Fixed candidate menus and evaluation units

Observed counts were thinned once per section or core into training (A), validation (B) and test (C) counts with fractions 0.5/0.25/0.25 by sequential binomial thinning[1]; windows and tiles take the rows of that one split, so the test molecules of a window never enter the section-wide loadings. Training counts constructed every geometry and fitted every component. Validation counts made every choice: density pooling, geometry, refinement penalty, scale, $\eta$, $\tau$, factor rank, gene-wise shrinkage where enabled, and the global and gene-wise mixture weights. Once every choice was fixed, the selected components were rebuilt on the combined training and validation counts with those settings unchanged and the per-bin normaliser recomputed (Methods, "Refit after selection"); test counts were never read, so held-out scores of the refit fields remain valid. The independently thinned-process interpretation is conditional on the working Poisson rates, not on the fixed observed count; given that count, split allocations are multinomial and dependent. The complete declared in-tissue mask, zero-count bins and original gene universe are shared by all candidates and all comparators.

| Item | Fixed specification |
|---|---|
| Fixed grids | Square cells of side $w \in \{1, 2, 4, 8, 16, 32\}$ bins, intersected with the support and split into four-connected components |
| Gaussian kernels | $\exp(-\|x_i - x_j\|^2/2\sigma^2)$ within $4\sigma$ bins, $\sigma \in \{0.5, 1, 2, 4, 8, 16\}$ bins, restricted to the support, not rescaled |
| Global endpoint | Spatially constant profile $q_g(\eta)$ |

| Item | Fixed specification |
|---|---|
| Joint Poisson merge | Greedy merge tree on the full training gene cube; snapshots at $K \in \{16, 32, \ldots, 4096\}$ regions with $K \leq N/4$ for $N$ support bins and $K$ at least the number of four-connected support components, which is the binding constraint on a ragged support; ties by region identifier |
| Boundary refinement | Objective $\sum_R F(R) - \lambda\,\text{per}$ with $\lambda \in \{3, 10, 30, 100\}$ nats per boundary edge; four checkerboard classes, each accepted or reverted as a whole; at most 20 sweeps; four refined versions of every snapshot enter the bank beside the unrefined snapshot |
| Global concentration | $\eta = 0.5$ throughout |
| Local concentration | $\tau \in \{1, 10, 10^2, 10^3, 10^4\}$, selected jointly with the geometry; whole-section tiled fits used $\{1, 10, \ldots, 10^5\}$ |
| Gene-wise shrinkage (optional) | $\tau_g \in \{1, 3, 10, \ldots, 10^4\}$ per gene, renormalised within each bin |
| Coarser sibling | Best candidate with at most $K/4$ regions (grid and adaptive families) or at least $4\sigma$ (Gaussian family); otherwise the coarsest candidate |
| Factor loadings | Training counts pooled into $4 \times 4$-bin units with at least 20 molecules over the whole section or core; Poisson non-negative matrix factorisation with multiplicative updates[2], three seeded restarts screened at 40 iterations and the best continued to 200; ranks $K \in \{8, 12, 20\}$; fitted once per section and shared by every window or tile |
| Factor weights | For every geometry in the bank: 30 multiplicative EM steps from uniform weights with a floor of $1/K$ per factor; one $\theta$ per region for regional geometries; winner and coarser sibling chosen on validation counts over all (geometry, rank) pairs |
| Factor-informed correction | Additive form, $(\sum_j W_{ij} Y_{Ajg} + \tau\, \widehat{\pi}_{ig}^{\mathrm{LR}})/(S_i + \tau)$, one component per selected spatial geometry, $\tau$ from the local concentration menu |
| Mixture weights | Global simplex weights by sequential least-squares programming[3]; gene-wise simplex weights with Dirichlet-type pull $\alpha = 1$ molecule per gene toward the global weights, 200 minorisation–maximisation iterations[4], renormalised within each bin |
| Density | Pooling selected on validation Poisson likelihood from the same bank; Gamma-regularised posterior mean (Methods) |
| Tiles | $128 \times 128$-bin squares at step 96 bins for supports above about $2 \times 10^4$ bins; tiles with fewer than 400 support bins dropped; every bin owned by the nearest centre among the tiles containing it, and a tile left owning no bin is dropped rather than fitted |

Here $F(R) = \sum_g C_{Rg} \log(C_{Rg}/|R|)$ with $0 \log 0 = 0$ is the joint Poisson profile score of a region, per the number of four-neighbour bin pairs with different labels, $W_{ij}$ the pooling weights of a geometry, $S_i = \sum_j W_{ij} T_{Aj}$, and $q_g(\eta)$ the global profile of Methods. Ties in the joint selection of geometry and $\tau$ favoured the global endpoint, then regional before Gaussian candidates, then fewer regions or larger bandwidth, and then smaller $\tau$. The component set of the gene-wise mixture is therefore the winner and coarser sibling of each of the grid, adaptive and Gaussian families, the factor winner and its sibling, and one factor-informed correction per selected spatial geometry. Native-bin size is retained throughout, so identical bandwidths in bins need not have identical physical dimensions.

The known-truth study used the same protocol on a wider geometry menu, as stated in Methods: the previous total-count guide, a greedy merge of total counts, the joint merge and the two-category target merge, with $\lambda \in \{0, 1, 3, 10\}$ and $K \in \{2, \ldots, 256\}$, and $\alpha$ chosen from $\{0, 1, 10, 100, 1000\}$ by a half split of the validation counts. Real data used the fixed $\alpha = 1$.

The evaluation units are as follows; paired comparisons count units on which one estimator exceeds another, and every unit is a technical or spatial replicate unless stated.

| Study | Units |
|---|---|
| Known truth | Four scenarios at 10 and 80 molecules per bin, 20 Poisson draws each; paired wins count draws |
| Same-section lung adenocarcinoma (Visium HD after Xenium) | Four non-overlapping $128 \times 128$-bin windows at 8 $\mu$m and thinning seeds 101–103: 12 window $\times$ seed units per experiment (289-gene panel; Xenium Prime 5K) |
| Renal cell carcinoma, Xenium Protein | Four $64 \times 64$-bin windows at 10 $\mu$m chosen on the PanCK image and seeds 101–103: 12 units |
| Non-small-cell lung cancer L1, post-Xenium immunofluorescence | Three $64 \times 64$-bin regions of interest at 10 $\mu$m chosen from the image and three seeds: 9 units; nine registration shifts of up to $\pm 10$ $\mu$m as sensitivity |
| Lung tissue microarray (Visium HD after Xenium) | Four cores from four donors, fitted whole-core in tiles, three seeds: 12 core $\times$ seed units, with donors reported alongside |
| Pulmonary fibrosis cohort (Xenium) | 45 cores, one thinning seed, fitted in tiles; a core contributes a unit for an annotation/programme pair only with at least one instance and 200 edge-zone bins; donor medians reported alongside; three cores refitted with two further seeds as sensitivity |

The Xenium reference of the same-section experiments, the protein channels, the immunofluorescence image and the pathologist polygons never entered fitting or selection. No test-count summary selected any model, and no claim of a globally optimal spatial partition follows from the finite search.

### Note 3. Programme definitions and biological interpretation

Every programme is a fixed list of panel genes, taken from panel membership or from the markers of the source paper and written down before any field was fitted; a programme probability is the sum of the gene probabilities over its genes (Methods). Genes absent from the shared gene set of a dataset were dropped from the programme, and a programme with no gene left was not scored. The references against which programmes are placed, the Xenium molecules of the same section, the protein and immunofluorescence channels and the pathologist polygons, never entered fitting or selection.

In the same-section lung adenocarcinoma experiments[5] the epithelial programme was EPCAM, CDH1 and MUC1, with NKX2-1 added in Experiment 2 where the panel carries it; the reference was the Xenium epithelial fraction of the same genes on the Visium HD grid. In the renal cell carcinoma Xenium Protein section[6] the endothelial programme (PECAM1, VWF) was compared with the CD31 channel and the epithelial programme (EPCAM, KRT8, KRT18, KRT7, CDH1, MUC1, KRT5) with the PanCK and E-cadherin channels; the two epithelial pairs are reported as uninformative in this tissue (Results). In the non-small-cell lung cancer section with post-Xenium immunofluorescence[7] the programme was EPCAM and KRT7; the public record does not state which antigen the imaged Opal 690 channel carries, so the compartment is described as an image compartment.

In the lung tissue microarray measured by Visium HD after Xenium[8], the programmes were fixed from the cohort design: the primary compartment programme was alveolar epithelium (AGER, RTKN2, SFTPC, NAPSA, SFTPD), with KRT17 (KRT17, ITGB6, MMP7) and fibrillar ECM (COL1A1, COL1A2, COL3A1, DCN, LUM) as secondary compartment programmes. All nine of the cohort's programmes were fitted whole-core on this slide and three were scored as compartment programmes; the other six have whole-core correlations only, where the core retained any of their genes above the abundance floor, because this slide carries no compartment reference for them, and the activated fibroblast programme was not a valid endpoint at all because the post-Xenium capture left it too few molecules. Cores were assigned to donors by matching the post-Xenium H&E to the cohort's per-core registered H&E, not by RNA (Methods).

In the pulmonary fibrosis cohort[8] nine programmes were fixed from the 343-gene panel before fitting: activated fibrotic fibroblast (CTHRC1, POSTN, FAP), fibrillar ECM (COL1A1, COL1A2, COL3A1, DCN, LUM), KRT17 associated epithelial (KRT17, ITGB6, MMP7), airway basal (KRT5, KRT15, TP63, KRT14), alveolar epithelium (AGER, RTKN2, SFTPC, NAPSA, SFTPD), airway secretory and ciliated (SCGB1A1, SCGB3A2, MUC5B, FOXJ1, C20orf85), endothelial (PECAM1, CLDN5, RAMP2, PLVAP, CA4), smooth muscle (ACTA2) and B-cell/TLS-associated (MS4A1, CD19, CD79A, CXCL13). These lists were prespecified for this analysis, informed by reported fibrotic fibroblast markers[8, 9], the $KRT5^{-}/KRT17^{+}$ epithelial state[8, 10] and lung compartments[11]. They are gene sums, not cell-type classifiers or signatures copied verbatim from those studies. In particular, the alveolar programme combines AT1- and AT2-associated transcripts, and the KRT17 score does not establish KRT5 negativity. Programme labels do not imply an exclusive cellular origin. The eight annotation/programme pairs were fibroblastic focus with the activated fibroblast programme; fibrosis, severe fibrosis and fibroblastic focus with fibrillar ECM; microscopic honeycombing and remodelled epithelium with airway basal; hyperplastic alveolar epithelium, and separately normal and minimally remodelled alveoli, with alveolar epithelium; large and small airway with airway secretory and ciliated; artery, muscularised artery and venule with smooth muscle; and tertiary lymphoid structure with the B-cell/TLS-associated programme. The annotations are the authors' own and were mapped into the Xenium frame from their cell-to-annotation table alone (Methods).

In cartilage[12] the programmes were cartilage matrix (COL2A1, ACAN, COL9A1) and collagen VI (COL6A1, COL6A2, COL6A3). These names describe gene panels; they do not assign measured RNA to extracellular protein compartments, and RNA on locations without a cell does not establish extracellular origin. The CODEX[13] analysis of the SPATCH pairs[14] uses no RNA programme: segmented CODEX cell density is a proxy for nuclear packing in a registered serial section, the epithelial fraction is the fraction of cells with the published epithelial annotation, and the 16 protein markers were prespecified from the panel. Serial sections do not measure identical cells, and nested regression increments are order-dependent and noncausal.

Observed group fractions pool molecules, whereas predicted programme summaries average bin probabilities; the two have different weighting and are labelled separately, and observed fractions of zero-total groups remain undefined. Mixture weights describe which geometry, scale or source predicts a gene; they are not cell or compartment proportions. Technical seeds, registration shifts and spatial windows are sensitivity analyses, not biological replication.

### Note 4. Development evidence and scope limits

The estimator was reached by single-variable changes, each tested with the protocol of Note 2 and kept only where validation-selected fields improved on an independent reference. The earlier released estimator built its adaptive regions from total counts alone and mixed one grid, one adaptive and one Gaussian component under a single global weight vector for all genes, with no gene-wise weights and no factor component. Its two fixed choices, the guide and the single weight vector, are ablated separately here: a total-count guide against the same merge on the full gene cube, and one global weight vector against gene-wise weights on today's geometry. The known-truth study states why both were replaced: geometry built from total counts cannot see a step of composition at constant density, whereas the same merge algorithm applied to the full gene cube can, and a global weight vector held the adaptive component low even for the genes it fitted best, whereas gene-wise weights let the programme genes take it (Results). The greedy merge of total counts is retained as the control that isolates the composition information in the partition, because the algorithm is otherwise identical. The earlier whole-section runs and the tile studies that preceded them are development evidence; their field-derived numbers are withdrawn from the main text and are not compared with the current fields.

Several alternatives were tested with the same protocol and not adopted. A multiplicative form of the factor-informed correction, the Poisson–Gamma posterior mean of a gene-specific rate multiplier around the factor prediction, gave more amplitude than an earlier additive configuration on the Xenium sections, but in the final configuration the additive prior overtakes it and it costs rank agreement on the 5,000-gene panel; the additive form is the default (Note 8). A logarithmic (geometric) pool of the components[15] in place of the per-bin renormalised linear mixture raised rank agreement slightly and lowered amplitude on the sparse panel; it was superseded by offering the factor component every geometry in the bank, which improved both. A hierarchical prior that shrinks the factor weights of a fine geometry toward those of its enclosing coarse geometry helped only while the factor component was confined to the geometries selected for the per-gene model, and was neutral once every geometry was offered; it remains a cheaper substitute when the candidate menu must be small. Shrinking each gene's mixture weights toward those of genes with similar loadings was neutral on both panels. Removing the plain spatial components so that only the factor component and its corrections remain beat the full set on every positional endpoint in 4 of 4 units of the 289-gene experiment, losing only the all-gene held-out score, and gave slightly more amplitude on the renal section; but, on the sixteen-gene truth where the factor prior is poor, lost edge accuracy and amplitude and fell below the target-tuned Gaussian in every structured scenario; the plain spatial components are the safety net when there is little to borrow across genes, and pruning is an option, not the default. Gene-wise shrinkage inside each component is likewise an option: it improved the all-gene held-out score in the known-truth study and restored part of the amplitude of the two sparse endothelial genes with spatial borrowing alone (Results), at some cost in rank agreement where the molecular source already supplies the amplitude; it is enabled where the Results say so. Two settings that are not alternatives but convergence matters were also fixed here: the gene-wise mixture must be run far enough that amplitude on the sparse panel has stopped rising, which the released 200 iterations achieve for the fields although not for the weights themselves (Note 9), and the factor loadings must be fitted on the whole section, because the units of one window are too few for a 5,000-gene panel (Methods). The sensitivity of the factor menu is in Supplementary Table 1: a rank menu, not a particular rank, is what matters, and unit size matters only when the units carry too few molecules.

The known-truth study also fixed two limits that must be read with the real-data results. First, held-out likelihood and edge accuracy are not the same objective, and at 80 molecules per bin the complete estimator gains on one while losing slightly on the other (Results). A factor prior built from sixteen genes is what leaves the two objectives apart; on real panels of hundreds to thousands of genes the positional references moved with the held-out score. Second, evidence bounds what any pooling can recover. In the sixteen-gene truth at 10 molecules per bin, spatial borrowing recovered less than half of the composition-only step and molecular borrowing had nothing to offer, so the factor component took 0.08 of the programme weight and no endpoint moved; a Gaussian tuned on the three target genes recovered 0.80 of the same step, and in a 300-gene panel the complete estimator recovered 0.73. What is recoverable is set by the programme's molecules in the window and the size of its compositional step, not by depth alone (Results). On the 5,000-gene panel the fields kept three quarters of the raw programme contrast where FICTURE kept more than the raw counts; that residual is shrinkage in the per-bin mixture, not an information bound, since the factor component alone reaches more of it (Results).

Held-out score gains of the same modality measure within-section prediction. They are not unbiased estimates or lower bounds of the mutual information between position and composition, and they are not evidence of a localisation mechanism; the positional claims rest on the independent references of Note 2. No comparison supports localisation below the bin scale: every reference compartment is itself an estimate, boundary distances are about one bin, and amounts are not comparable between chemistries measured on the same section. Windows, regions of interest and cores are technical or spatial units; only the fibrosis cohort offers donors as replication units. Stitched whole-section fields can carry seams at tile boundaries, and no smoothing across seams was applied before any endpoint.

### Note 5. Captured RNA density contains information beyond cell packing

We asked whether captured RNA per area could be summarised by cell packing alone. In registered SPATCH colon adenocarcinoma (COAD), hepatocellular carcinoma (HCC) and ovarian cancer (OV) sections[14], we calculated observed UMI totals per represented tissue area in common 200-$\mu$m units, with CODEX segmented-cell density as the packing proxy. On spatially held-out blocks, the cell-density model achieved out-of-fold $R^2$ values of 0.257, 0.110 and 0.144; adding epithelial fraction and the 16 measured protein markers increased these to 0.500, 0.420 and 0.405 on the same units (Extended Data Fig. 6; Supplementary Tables 2–5). Epithelial fraction alone improved prediction in COAD and HCC but not OV ($R^2 = 0.321$, 0.225 and 0.143). Captured density therefore carries molecular information beyond the segmented-cell count. This analysis uses observed counts, not fitted fields; because the protein panel includes lineage markers and the measurements come from registered serial sections, it does not isolate a cell-intrinsic state effect or establish causality. Each cancer is one supplied specimen pair rather than a cancer-level replication, and the intervals are within-section.

### Note 6. Robustness of thinning-based selection to overdispersion

Binomial thinning yields independent Poisson splits only for Poisson counts; under a negative-binomial count with mean $\mu$ and dispersion $\phi$ (variance $\mu + \mu^2/\phi$) the thinned training and validation counts of the same bin and gene are positively correlated, with covariance $f_A f_B \mu^2/\phi$, so that validation could reward pooling that reproduces noise shared with training. We quantified this on the 300-gene known-truth simulation (Extended Data Fig. 2) by drawing counts as negative binomial with the same means and $\phi \in \{\infty, 10, 3, 1\}$, independently per bin and gene, and applying the unchanged pipeline (20 draws per setting). At 10 molecules per bin the programme edge error and the background root mean squared error moved by at most 0.002 at any $\phi$ and the recovered contrast by no more than one standard error, the validation split and an independent validation draw selected candidates of the same true accuracy, and the validation score ranked the geometry-by-$\tau$ menu against the truth as under Poisson (Spearman 0.80–0.86). At 80 molecules per bin, with the dispersion applied independently per bin and gene, the paired ordering of estimators was preserved in 20 of 20 draws at every $\phi$ (edge error, contrast, background and all-gene held-out score), the selected shrinkage and the deep-gene calibration were unchanged for $\phi \geq 3$, and the empirical split correlations matched the closed form (0.10 for the deepest genes at $\phi = 3$). Only at $\phi = 1$, where the deepest genes carry about one molecule per bin with geometric variance and a split correlation of 0.24, did validation select native-bin geometries in every draw whereas an independent validation draw selected the oracle's geometry; the complete estimator then still beat the spatial-only fields on every endpoint but recovered 0.73 rather than 0.96 of the true contrast and shrank the 20 deepest genes to a slope of 0.56 on the truth.

Two results run the other way and bound the claim. First, the estimator's own all-gene selection is more robust than a selection made on the programme alone: at 80 molecules per bin and $\phi = 3$ a target-only criterion picked a candidate worse than an independent draw's in 12 of 20 draws, moving to a $\sigma$ of 1 or 0.5 bins, while the all-gene criterion used by the package matched the oracle's pick with a regret of 0.0000 and ranked the menu at Spearman 0.995. Second, a multiplier shared by all genes of a bin (bin-level overdispersion, split correlation 0.90–0.96 of the totals at 80 molecules per bin) left every selection unchanged, because composition is multinomial conditional on the bin total, but it does add edge error to all three estimators: at $\phi = 3$ the complete estimator's edge error rose from 0.0053 to 0.0158 and the spatial-only stack's from 0.0164 to 0.0227, and at $\phi = 1$ the complete estimator's edge advantage over spatial borrowing was gone (6 of 20 draws, mean 0.0256 against 0.0249), though contrast, background and the all-gene score still favoured it in 18 to 20 of 20. Selection is therefore robust to bin-level overdispersion while edge placement is not. At per-gene means above about $\phi$ molecules per bin, a separately generated validation draw would avoid the shared count fluctuation responsible for the observed selection failure; another thinning of the same counts would not. Source: research/nb_robustness_20260909.

### Note 7. What the simulations do and do not reproduce

The known-truth simulations establish which design choices are necessary and when the molecular source is used, and they are deliberately simpler than a tissue panel. Three consequences follow and are stated here rather than in the Results. First, when the simulated truth is exactly low rank a gene's own pooled counts cannot improve on the factor field in likelihood, so the weight a gene places on the raw factor component rises with its depth; introducing gene-specific departures from the low-rank truth restores weight to the corrected components at 80 molecules per bin but not at 10, and the allocation measured on real panels (Fig. 3d of the main text) is reported as measured rather than as reproduced by a simulation. Second, the factor field's geometry is chosen once on the evidence of all genes, so in simulations whose background varies while the target does not, the target's edge is placed less well even as the held-out score and the amplitude improve; this happens in the structured background, in the truth with no co-expressed members and in the independent-gene truth, and offering the factor field on every selected geometry halves that loss at 10 molecules per bin and reduces it by a fifth at 80. Third, a factor source that carries no information is not left at zero weight. On an independent-gene truth, where no gene follows the programme's step, the target genes still placed 0.32 of their weight on the factor components at 10 molecules per bin and 0.71 at 80 (Extended Data Fig. 2b). What that source is worth there is nothing at 10 molecules per bin, where no endpoint changes, and a small trade at 80, where a little edge accuracy is exchanged for a little contrast. The claim the design supports is that an uninformative source changes no endpoint, not that it receives no weight. Held-out likelihood and edge accuracy are distinct objectives, as they are for any selection rule, and the manuscript reports both throughout rather than optimising one against the other. Source: research/synthetic_factor_20260909.

### Note 8. Alternatives implemented and not adopted

Two variants of the factor-informed correction were implemented and measured. A multiplicative form, the Poisson–Gamma posterior mean of a gene-specific rate multiplier around the factor prediction, recovered more programme amplitude than an earlier additive configuration on the two Xenium sections, but in the final configuration the additive prior overtakes it, beating it on every endpoint of the immunofluorescence section in a paired comparison, and it costs rank agreement on the 5,000-gene panel (0.446 against 0.470, in 0 of 4 windows); the additive form is used throughout. A per-gene shrinkage inside the corrected components raised each correction's own validation likelihood by 0.05, 0.009 and 0.008 nats per validation molecule on the 289-gene, 5,000-gene and renal panels, and the shrinkage it selected fell with a gene's depth as expected; once the mixture was refitted, however, weight moved onto the corrected components and left the plain spatial components nearly empty (0.11 to 0.01 of the weight on the 5,000-gene panel), which cost rank agreement there, so a single shrinkage per component is kept. That the decision then falls to the mixture is not an escape: Note 9 reaches the same conclusion from the other side, by offering each geometry its correction at every shrinkage on the menu so that a gene selects its own effective shrinkage through the mixture weights rather than within a component, which moves the corrected class to 0.91 of the weight and costs 0.037 of edge rank agreement on the 5,000-gene panel while gaining 0.005 nats on the 289-gene panel. Per-gene shrinkage toward the factor field is therefore not a matter of where the choice is made. On the immunofluorescence section the option is unambiguously better, winning 9 of 9 units on contrast, band error, the zone held-out binomial and the all-gene held-out score, so what it costs is rank agreement on a dense panel and what it buys is calibration on a sparse one. A third option offers the factor field on every selected spatial geometry rather than on the all-gene winner and its coarse sibling alone; its effect on a structured-background simulation is the yellow series of Extended Data Fig. 2 and is described in Note 7. A fourth, a compact form that keeps only the factor component and the corrected spatial components and drops the plain spatial ones, is left off because on small panels the factor prior is poor and the plain components are what the mixture falls back on. All four are available in the package and off by default. Source: research/genewise_tau_20260909, research/lowrank_candidate_20260908 and research/synthetic_factor_20260909.

### Note 9. What the per-gene weights are and are not

The mixture assigns every gene a simplex weight over its selected components, and those weights are a fitting device rather than a measurement of where the gene borrows. We fixed every choice made on validation counts for a window — geometries, $\eta$, spatial and correction shrinkages, factor rank — and refitted only the weights from nine starting points, the package's own, a uniform start, four Dirichlet draws and three adversarial vertices, at three penalty strengths.

The delivered field is stable and the split between classes is not. Across those starts the mean absolute difference in the combined composition field was at most $2.1 \times 10^{-4}$ and the Spearman correlation of the programme fraction at least 0.987, while the mean weight on the corrected class moved by up to 0.16 on the 289-gene panel and 0.44 on the 5,000-gene panel at the released 200 iterations. Run to 2,000 iterations the class spans fall below 0.05, so the objective has one optimum and is nearly flat around it; the released setting stops at a point whose weights differ while its field does not. Running the weights to convergence moves no reported endpoint beyond the third decimal and does not move them consistently: on the 289-gene panel only the all-gene held-out score improves in all four windows, by 0.0013 nats, while each positional endpoint improves in two or three of four, and on the 5,000-gene panel the held-out score is worse in all four, so stopping at 200 iterations acts there as implicit regularisation of a weight fit made on validation counts. Per gene the movement is larger: at the settings used here a quarter of genes can have their corrected-class weight moved by more than 0.06 to 0.12, and the most movable gene can be turned from almost all corrected to almost all factor. Independently, the same gene's factor weight differs by a median of 0.25 to 0.33 between adjacent windows of one section.

The shrinkage menu moves the split as much as the starting point does. Widening the $\tau$ menu from the released five values to seven leaves the held-out likelihood and every positional endpoint unchanged to three decimal places while doubling the weight on plain spatial pooling, because the released menu is truncated: on the 5,000-gene panel four of six geometries select its upper value. Offering a gene a menu of shrinkages on one geometry, rather than the single selected one, moves the corrected class to 0.91 of the weight; that variant is not among the package's options, since one corrected component per geometry enters the mixture, and its measurements are in Note 8.

Two things are determined and are worth stating. The correction's own shrinkage is sharply identified on validation counts, with the extremes of the menu losing thousands of nats, so a corrected component is a well-defined object rather than a redundant reparametrisation of its two limits. And a leave-one-class-out ablation with every other choice held fixed, run to convergence, shows the two sources are jointly necessary and that their balance differs by panel: with 289 genes the corrected class buys held-out likelihood and amplitude at a small cost in placement, whereas with 5,000 genes the corrected class alone reproduces the complete estimator on every endpoint and the remaining classes add 0.001 nats. Source: research/mixture_identifiability_20260909.

#### Supplementary Table 1. Sensitivity of the factor menu

Same-section Experiment 2 (5,000-gene panel), four 1-mm windows, one thinning seed; the complete estimator with section-wide loadings fitted on units of the stated size for the stated rank menu. Edge-zone Spearman correlation and AUC against the Xenium epithelial fraction, area-matched overlap, boundary distance (bins), all-gene held-out score (nats per molecule) and wall time per window on eight threads. Source: research/lowrank_candidate_20260908/sens_exp2.

| Rank menu | Unit (bins) | Spearman | AUC | IoU | Boundary | Held-out | Minutes |
|---|---|---|---|---|---|---|---|
| 8, 12, 20 (default) | 4 | 0.469 | 0.821 | 0.742 | 1.09 | −7.344 | 15 |
| 4, 8, 12, 20, 30 | 4 | 0.469 | 0.821 | 0.742 | 1.09 | −7.344 | 20 |
| 12 only | 4 | 0.404 | 0.784 | 0.711 | 1.37 | −7.348 | 9 |
| 8, 12, 20 | 2 | 0.425 | 0.800 | 0.722 | 1.32 | −7.354 | 21 |
| 8, 12, 20 | 8 | 0.472 | 0.825 | 0.749 | 1.06 | −7.341 | 11 |

With the wider menu validation chose the same rank (8) in every window, so the extra ranks were unused; a single rank is what costs agreement.

### Supplementary Tables 2–5. CODEX native-count analysis

Supplementary Tables 2–5 support the captured-density analysis of the registered SPATCH sections (Note 5 and Discussion) and are independent of the estimator. They use the common-bin native-count analysis for COAD, HCC and OV separately. The response is exact UMI divided by represented area. Packing, epithelial fraction and all 16 fixed marker means are fitted on the same eligible units. OOF denotes held-out 1-mm-block fold predictions; folds are spatially blocked[16] but have no exclusion buffer between training and test blocks. Thus OOF performance is a within-section prediction sensitivity rather than independent-section validation. Intervals are 500 within-section block-bootstrap percentile intervals[17], not patient-level confidence intervals. Negative OOF values, where present, are retained. Individual-marker partial correlations adjust for packing and epithelial fraction (Supplementary Table 5); the corresponding increments are fitted $R^2$ changes after those controls (Supplementary Table 4). The out-of-fold $R^2$ values quoted in Results are the raw-scale rows of Supplementary Table 3; the eligible unit counts in the figure legend are those of Supplementary Table 2.

#### Supplementary Table 2. CODEX paired-area sample units

| Pair | RNA bins | Zero bins | CODEX cells | Eligible units | 1-mm blocks |
|---|---|---|---|---|---|
| COAD | 650209 | 31 | 193782 | 1063 | 49 |
| HCC | 671560 | 0 | 69076 | 1095 | 48 |
| OV | 629672 | 98 | 166186 | 1027 | 47 |

#### Supplementary Table 3. CODEX density models

| Pair | Scale | Model | Fitted R2 | OOF R2 | Fitted R2 interval |
|---|---|---|---|---|---|
| COAD | raw | Packing | 0.2614 | 0.2569 | 0.1645, 0.3766 |
| COAD | raw | + epithelial | 0.3265 | 0.3208 | 0.2427, 0.4185 |
| COAD | raw | + 16 markers | 0.5840 | 0.5002 | 0.5321, 0.6779 |
| COAD | log1p | Packing | 0.3135 | 0.3102 | 0.2159, 0.4219 |
| COAD | log1p | + epithelial | 0.3647 | 0.3610 | 0.2806, 0.4522 |
| COAD | log1p | + 16 markers | 0.5793 | 0.4794 | 0.5279, 0.6967 |

| Pair | Scale | Model | Fitted R2 | OOF R2 | Fitted R2 interval |
|---|---|---|---|---|---|
| HCC | raw | Packing | 0.1155 | 0.1103 | 0.0264, 0.2357 |
| HCC | raw | + epithelial | 0.2324 | 0.2247 | 0.0990, 0.3929 |
| HCC | raw | + 16 markers | 0.4827 | 0.4197 | 0.3596, 0.6436 |
| HCC | log1p | Packing | 0.0888 | 0.0852 | 0.0125, 0.2088 |
| HCC | log1p | + epithelial | 0.1892 | 0.1827 | 0.0696, 0.3548 |
| HCC | log1p | + 16 markers | 0.4442 | 0.3822 | 0.3420, 0.6151 |
| OV | raw | Packing | 0.1631 | 0.1445 | 0.0859, 0.2943 |
| OV | raw | + epithelial | 0.1702 | 0.1425 | 0.1023, 0.2963 |
| OV | raw | + 16 markers | 0.4678 | 0.4048 | 0.3900, 0.6710 |
| OV | log1p | Packing | 0.2398 | 0.2250 | 0.1546, 0.3352 |
| OV | log1p | + epithelial | 0.2413 | 0.2150 | 0.1552, 0.3481 |
| OV | log1p | + 16 markers | 0.6764 | 0.6205 | 0.6171, 0.7780 |

### Supplementary Table 4. CODEX nested fitted increments

| Pair | Scale | Addition | R2 increment | Interval |
|---|---|---|---|---|
| COAD | raw | epithelial | 0.0650 | 0.0214, 0.1247 |
| COAD | raw | marker | 0.2575 | 0.1756, 0.3888 |
| COAD | log1p | epithelial | 0.0512 | 0.0092, 0.1181 |
| COAD | log1p | marker | 0.2146 | 0.1509, 0.3391 |
| HCC | raw | epithelial | 0.1169 | 0.0533, 0.1974 |
| HCC | raw | marker | 0.2503 | 0.1821, 0.3595 |
| HCC | log1p | epithelial | 0.1005 | 0.0393, 0.1823 |
| HCC | log1p | marker | 0.2549 | 0.1825, 0.3736 |
| OV | raw | epithelial | 0.0071 | 0.0000, 0.0404 |
| OV | raw | marker | 0.2977 | 0.2311, 0.4404 |
| OV | log1p | epithelial | 0.0015 | 0.0000, 0.0286 |
| OV | log1p | marker | 0.4351 | 0.3541, 0.5571 |

### Supplementary Table 5. Individual CODEX markers

| Pair | Marker | Partial correlation | R2 increment |
|---|---|---|---|
| COAD | CD8 | -0.2206 | 0.0328 |
| COAD | CD20 | 0.4139 | 0.1154 |
| COAD | CD3e | 0.0333 | 0.0007 |
| COAD | CD56 | -0.3122 | 0.0657 |
| COAD | Pan-Cytokeratin | -0.2156 | 0.0313 |
| COAD | CD4 | 0.0594 | 0.0024 |
| COAD | CD34 | -0.2428 | 0.0397 |
| COAD | SMA | -0.1011 | 0.0069 |
| COAD | FOXP3 | -0.1716 | 0.0198 |
| COAD | CD163 | -0.2624 | 0.0464 |
| COAD | HLA-A | -0.1626 | 0.0178 |
| COAD | CD11c | -0.2012 | 0.0273 |
| COAD | MPO | -0.2223 | 0.0333 |
| COAD | CD68 | -0.2704 | 0.0492 |
| COAD | HLA-DR | -0.0424 | 0.0012 |
| COAD | IDO1 | 0.0300 | 0.0006 |
| HCC | CD8 | 0.0137 | 0.0001 |
| HCC | CD20 | 0.2289 | 0.0402 |
| HCC | CD3e | 0.0593 | 0.0027 |
| HCC | CD56 | -0.1333 | 0.0136 |
| HCC | Pan-Cytokeratin | 0.2486 | 0.0474 |
| HCC | CD4 | 0.2324 | 0.0415 |
| HCC | CD34 | 0.0496 | 0.0019 |
| HCC | SMA | -0.1828 | 0.0256 |
| HCC | FOXP3 | 0.0187 | 0.0003 |

| Pair | Marker | Partial correlation | R2 increment |
|---|---|---|---|
| HCC | CD163 | 0.0508 | 0.0020 |
| HCC | HLA-A | 0.1301 | 0.0130 |
| HCC | CD11c | 0.2330 | 0.0417 |
| HCC | MPO | -0.0595 | 0.0027 |
| HCC | CD68 | 0.1587 | 0.0193 |
| HCC | HLA-DR | 0.2104 | 0.0340 |
| HCC | IDO1 | 0.0730 | 0.0041 |
| OV | CD8 | 0.3476 | 0.1003 |
| OV | CD20 | 0.0759 | 0.0048 |
| OV | CD3e | 0.2816 | 0.0658 |
| OV | CD56 | 0.4433 | 0.1630 |
| OV | Pan-Cytokeratin | -0.1716 | 0.0244 |
| OV | CD4 | 0.3288 | 0.0897 |
| OV | CD34 | 0.2838 | 0.0668 |
| OV | SMA | -0.0548 | 0.0025 |
| OV | FOXP3 | 0.2963 | 0.0729 |
| OV | CD163 | 0.2139 | 0.0380 |
| OV | HLA-A | -0.1181 | 0.0116 |
| OV | CD11c | 0.3969 | 0.1307 |
| OV | MPO | -0.0378 | 0.0012 |
| OV | CD68 | -0.1768 | 0.0259 |
| OV | HLA-DR | 0.0678 | 0.0038 |
| OV | IDO1 | 0.0541 | 0.0024 |
